# Transcriptome and epigenome dynamics underpin cold stress priming in *Arabidopsis*

**DOI:** 10.64898/2025.12.16.694799

**Authors:** Munissa Sadykova, Hidetoshi Saze

**Affiliations:** Plant Epigenetics Unit, Okinawa Institute of Science and Technology Graduate University (OIST), Onna-son, Okinawa, 904-0495, Japan

## Abstract

Stress priming is a critical adaptive mechanism that enables plants to enhance responses to recurring environmental stresses. While transcriptomic changes associated with cold stress priming have been reported, the underlying epigenetic mechanisms remain largely unknown. Here, we investigated transcriptomic and DNA methylation dynamics during cold priming in *Arabidopsis thaliana*. Cold stress induces distinct gene expression patterns between primed and non-primed plants, accompanied by DNA methylation changes across all cytosine contexts in both protein-coding genes and transposable elements (TEs). Furthermore, DNA methylation mutants exhibit altered cold stress memory, highlighting a role for DNA methylation in preventing spurious gene activation and maintaining priming specificity. Notably, *met1*, deficient in CG methylation, shows reduced methylation at the *CBF* gene cluster, which correlates with its overexpression and enhanced activation of downstream cold-responsive genes. Our findings demonstrate that DNA methylation modulates cold stress memory by shaping chromatin states and ensuring transcriptional precision.

## Introduction

To endure fluctuating environmental conditions, such as extreme temperatures, plants rely on robust stress response mechanisms. Cold stress, which includes both chilling (0 – 14 °C) and freezing (< 0 °C) temperatures, has significant impacts on plant growth, development, and crop productivity ^1^. To cope with cold stress, plants initiate a cascade of responses that facilitate cold acclimation upon sensing temperature changes at the cell membrane – a process through which plants acquire higher tolerance to freezing conditions upon exposure to milder, non-freezing temperatures ^2^.

In *Arabidopsis thaliana* and other plant species, CBF/DREB1 (C-REPEAT BINDING FACTORS/DEHYDRATION RESPONSIVE ELEMENT-BINDING 1) transcription factors (TFs) orchestrate transcriptomic changes during cold acclimation ^3–5^. In the *A. thaliana* genome, *CBF1*, *CBF2*, and *CBF3* are in a tandem cluster, and are rapidly activated under cold stress by ICE1 (INDUCER OF CBF-EXPRESSION 1) and other TFs, which in turn induce the expression of downstream *COR* (COLD-REGULATED) genes ^6,7^. *COR* genes encode chaperones, dehydrins, and cryoprotective proteins that help protect plant cells from freezing damage by stabilizing chloroplast and plasma membranes ^8–11^.

Beyond immediate stress responses, plants that are exposed to a priming stress stimulus can acquire higher tolerance to subsequent stresses, indicating the presence of a mechanism that allows plants to retain memory of previous exposure to stress, and to adapt to recurrent stressors^12^. In *A. thaliana*, this acquired freezing tolerance in primed plants is accompanied by distinct transcriptomic and metabolomic responses ^13^. Such transcriptional memory is characterized by either enhanced or attenuated gene expression, or sensitization of genes to stress cues, and are often associated with changes in chromatin state, including DNA methylation, histone modifications, and production of small interfering RNA (siRNA) and other non-coding RNAs (ncRNAs) ^14^. These chromatin modifications are often interdependent, and are critical for regulating gene expression, maintaining genome integrity, and silencing transposable elements (TEs) ^15,16^.

In plants, DNA methylation is established *de novo* through the plant-specific RNA-directed DNA methylation (RdDM) pathway ^17,18^. In this pathway, RNA polymerase IV (Pol IV) is recruited to RdDM loci by the histone-binding protein SHH1 (SAWADEE HOMEODOMAIN HOMOLOG 1), which recognizes H3K9me2 marks ^16^. The single-stranded RNAs produced by Pol IV are then converted into double-stranded RNAs (dsRNAs) by RDR2 (RNA-DEPENDENT RNA POLYMERASE 2) which are then processed into 24-nt siRNAs by DCL3 (DICER-LIKE 3) ^19–21^. These siRNAs are loaded into ARGONAUTE proteins within the RNA-induced silencing complex (RISC), which guides the DNA methyltransferase DRM2 (DOMAINS REARRANGED METHYLTRANSFERASE 2) to target loci, enabling methylation of cytosines in CG, CHG, and CHH (where H = A, C, or T) sequence contexts ^22^. Once established, DNA methylation is maintained in the CG context by MET1 (METHYLTRANSFERASE 1), a member of the DNMT1 (DNA (CYTOSINE-5)-METHYLTRANSFERASE 1) family ^23^. Plant-specific CMT3 (CHROMOMETHYLASE 3) and CMT2 bind H3K9me and mediate methylation in CHG and CHH contexts, respectively ^18,24–26^. Methylation in heterochromatic regions is further supported by the chromatin remodeler DDM1 (DECREASED DNA METHYLATION 1), which facilitates access of methyltransferases to nucleosome-bound DNA ^26,27^.

The importance of epigenetic regulation of chromatin has been demonstrated under various abiotic stress conditions, including drought, salinity, heat, and cold ^28–35^. For example, in rice and maize, cold stress induces accumulation of histone acetylation at the promoters of cold-responsive genes, including *CBF/DREB1*. In *A. thaliana*, increased histone acetylation has been observed at *COR* genes, such as *COR15A* and *COR47* ^36^. Furthermore, cold treatment leads to rapid redistribution of the antagonistic histone 3 lysine 4 trimethylation (H3K4me3) and H3K27me3 marks at *CBF/DREB1* and *COR* genes in *A. thaliana* ^35,37^. However, these cold-induced epigenetic changes are not always associated with alteration of gene transcription ^35,37^, and it remains unclear how epigenetic mechanisms regulate cold stress responses, priming, and stress memory formation in plants.

In this study, we investigated the role of DNA methylation in cold stress priming and memory in *A. thaliana* using wild-type (WT) and DNA methylation-deficient mutants. Our findings show that cold stress induces significant DNA hypomethylation in protein-coding genes and TE regions, particularly in the CHH context. We identified genes that display either enhanced or attenuated gene expression after cold priming and repeated cold exposure. These cold stress memory genes, which include TEs and ncRNAs, exhibit low DNA methylation levels and are significantly enriched in active chromatin states. Intriguingly, most cold memory genes identified in WT background were not primed in DNA methylation mutants. Instead, over a thousand new memory genes were primed in these mutants, suggesting that DNA methylation shapes the specificity of transcriptional memory under cold stress. Notably, we observed strong upregulation of *CBF/DREB1* genes in *met1* mutants, highlighting the role of CG methylation in cold acclimation. Together, our findings demonstrate that DNA methylation is essential for regulating transcriptional responses to cold stress and establishing cold stress memory in *A. thaliana*.

## Results

### Identification of cold stress memory genes in *A. thaliana*

To investigate the impact of cold priming and stress memory on global transcriptomic dynamics, we performed transcriptome analysis of cold-primed and non-primed *Arabidopsis* plants as illustrated in Fig. 1a. *Arabidopsis* WT Col-0 plants were grown in a standard condition (see ‘Materials and Methods’) at 22°C for 12 days and exposed to the first (priming) cold treatment at 4°C for 3 days. The plants were further placed in the standard condition for a lag (recovery) phase for 7 days and exposed to the second (triggering) cold treatment. The transcriptome of plants with no cold treatment (control; C) was compared with plants exposed to priming cold treatment only (priming; P), triggering cold treatment only (triggering; T), and the priming and triggering condition (Priming and Triggering; PT) by RNA-seq analysis with three biological replicates (Fig. S1a).

**Figure 1.**
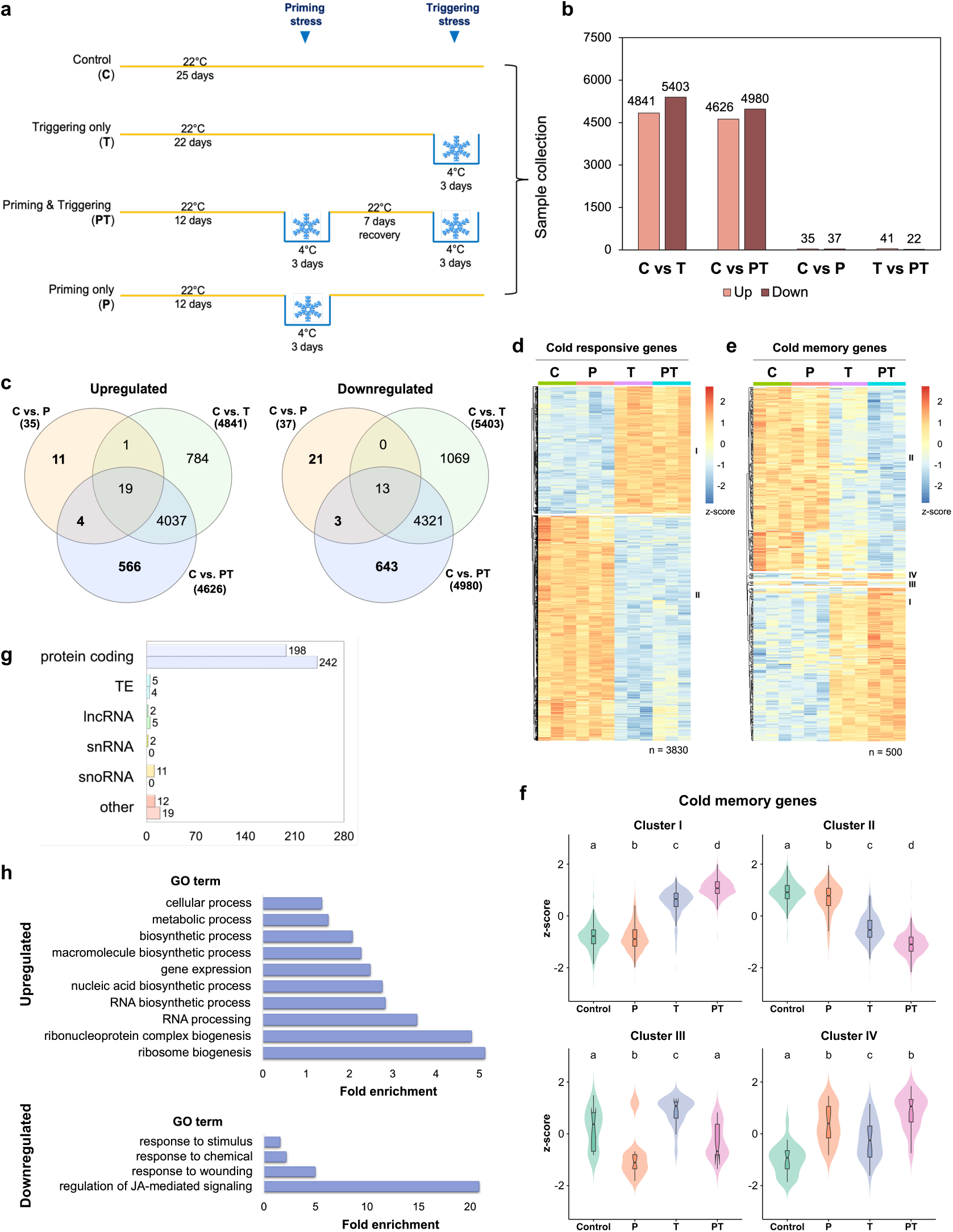
Cold stress priming alters transcriptomic response in *A. thaliana*. **a**. Experimental setup for the cold stress treatment of plants. Plants were treated with Priming only (P), Triggering only (T), Priming and Triggering (PT), or no cold stress treatment (Control, C). For the stress treatment, plants were moved to 4°C for 3 days. The recovery period between priming and triggering stress lasted for 7 days at 22°C. All samples were collected at 25-days-post-germination. **b**. Numbers of differentially expressed genes (DEGs) between the Control and cold-treated groups (*p-adj* < 0.05) in wild-type Col-0 (WT) *A. thaliana*. C: Control, T: Triggering only, PT: Priming and Triggering, P: Priming only. **c**. Venn diagrams between the upregulated and downregulated DEGs in Control vs. P, Control vs. T, and Control vs. PT comparisons. Genes identified as cold stress memory are highlighted in bold. **d**, **e**. Heatmap of expression of cold-responsive genes (d) and cold memory genes (e) in WT plants. DEGs with *p-adj* < 0.05 and |log2FoldChange (FC)| > 0.585 were selected. Gene expression values (log2FC) were scaled by row to obtain z-scores and rows were clustered using Euclidian distance. **f**. Violin plots of mean genes expression of cold stress memory genes by clusters. Statistical significance was determined using pairwise Wilcoxon rank-sum tests. P-values were adjusted using Benjamini-Hochberg (BH) method. Different letters correspond to significant differences between the treatment groups (*p-adj* < 0.01). **g**. Gene categories among the WT cold memory genes (n = 500). Top bar: genes upregulated in PT; bottom bar: genes downregulated in PT plants. pre-tRNA, ribosomal RNA, miRNA primary transcript, pseudogene, and novel transcribed region were classified as ‘other’. **h**. GO term analysis of WT cold memory genes performed using PANTHER. Fold enrichment of GO terms with Bonferroni-corrected P-values < 0.05 are displayed.

We identified 10,244 differentially expressed genes (DEGs) between C and the T plants, with 4,841 genes significantly upregulated and 5,403 genes downregulated in the T group (Fig. 1b, Supplementary Data 1). In the PT group, 9,616 DEGs were identified compared to the C, with 4,626 genes upregulated and 4,980 genes downregulated in the PT (Fig. 1b). This contrasts with the 72 DEGs found between C and P plants, suggesting that most cold-induced gene expression returns to baseline levels within 10 days post-treatment (Fig. 1b, Fig. S1a).

Stress priming can result in enhanced up- or down-regulation of genes upon exposure to a second stimulus, as well as sensitization of genes to the stress signal or prolonged induction ^14,38^. In this study, we classified DEGs between C and T as cold-responsive genes, while remaining DEGs between C and P, C and PT, and T and PT were classified as cold stress memory genes (memory genes) whose expression profile were influenced by the presence or absence of the priming treatment (Fig. 1c). As a result, 3,830 cold-responsive genes were identified between C and T conditions (Fig. 1d, Fig. S1b, Supplementary Data 1), which showed largely consistent responses to both the first and second cold treatments. In contrast, a total of 500 genes were identified as cold memory genes (Fig. 1e–f, Supplementary Data 1), showing altered transcriptional responses to repeated cold stress. More than 90% of both cold responsive and memory genes were protein-coding genes (Fig. S1c, Fig. 1g).

Gene Ontology (GO) analysis of the cold-responsive genes revealed that genes related to cold response and acclimation were upregulated, whereas downregulated genes were enriched for terms related to photosynthesis and metabolic processes, such as glucosinolate and indolalkylamine biosynthesis (Fig. S1d, Supplementary Table 1). In turn, genes upregulated in the primed plants were enriched for metabolic processes and nucleic acid synthesis, while downregulated genes were enriched for terms such as response to stimulus and JA-mediated signalling regulation (Fig. 1h, Supplementary Table 1). We found that rRNA processing and ribosome biogenesis processes were upregulated in both primed and non-primed plants (Fig. 1h, S1d), confirming their importance in cold tolerance of the plants ^39–41^.

In addition to protein-coding genes, we found 20 TEs and 27 ncRNAs, including the lncRNA *SVALKA* involved in the transcriptional regulation of *CBF* genes ^42,43^, as cold-responsive genes (Fig. S1c–e). Additionally, we identified 9 TEs and 20 ncRNAs, including snRNAs and snoRNAs, as cold memory genes (Fig. 1g, Fig. S1f–g). The cold responsive TEs were enriched for LTR (Long Terminal Repeat) retrotransposons from *Gypsy* and *Copia* superfamilies (Fig. S2a), including the *ATCOPIA46* (Fig. S2b–c), which shows a sequence similarity to a heat-responsive retrotransposon family in *Brassicaceae* ^44^. These results suggested a broad impact of cold stress response and priming on transcriptional regulation of both protein-coding and non-coding genes in *Arabidopsis*.

### Cold stress priming results in global DNA methylation changes

Previous studies of DNA methylation in plants have suggested its role in acquiring stress tolerance and adapting to changing environments ^45–48^. To understand the dynamics of DNA methylation during cold stress response and memory regulation in *A. thaliana*, we performed global DNA methylation analysis using EM-seq on plants subjected to C, P, T, and PT treatments (Fig. 1a). The results revealed a slight increase in CG and CHG methylation in the bodies of protein-coding genes under the P condition, while significant hypomethylation – particularly in CHH contexts – under the T and PT conditions (Fig. 2a). Similar to protein-coding genes, TEs showed elevated CG and CHG methylation under the P condition, but were significantly hypomethylated in the CHH context under the PT condition (Fig. 2b).

**Figure 2.**
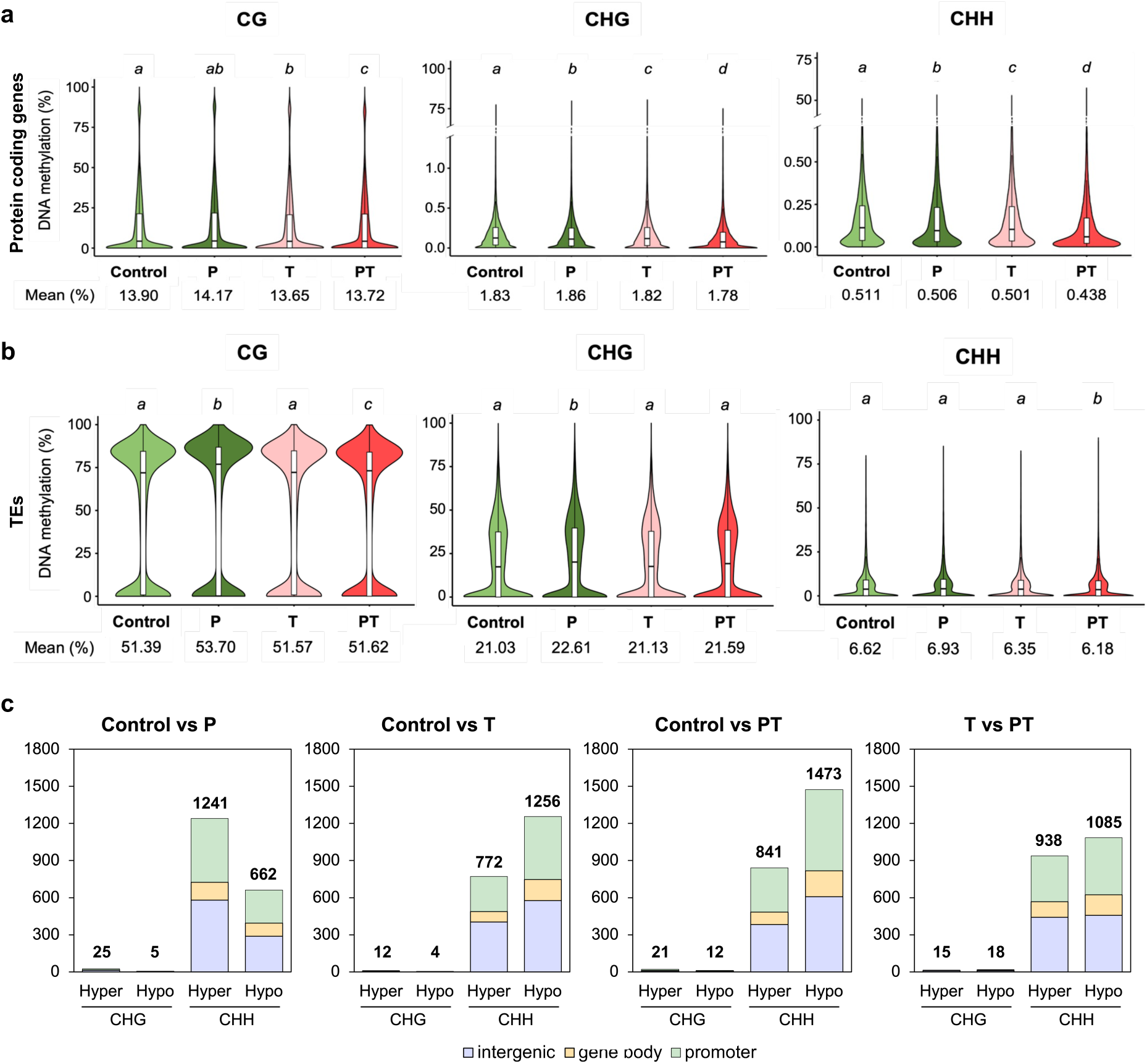
Cold stress and priming result in global DNA hypomethylation in *A. thaliana*. **a, b**. Violin plots of mean DNA methylation in the gene body of protein-coding genes (a) and TEs (b). Statistical significance was assessed using pairwise Wilcoxon rank-sum tests. Different letters indicate significant differences between treatment groups (*p-adj* < 0.05). **c**. Distribution of DMRs in intergenic regions, gene body, and promoter regions. DMRs were identified using a minimum methylation difference threshold of 40% for the CG context, 25% for the CHG, and 10% for the CHH contexts. The minimum overlap required for DMRs was set to 1 bp. Promoter regions were arbitrarily selected as 1 kb region upstream of genes.

To identify regions undergoing significant changes in DNA methylation, we extracted differentially methylated regions (DMRs) in the CG, CHG, and CHH contexts. The results revealed about 2,000 DMRs between control and cold-treated plants (Fig. 2c, Supplementary Data 2). Notably, a large number of DMRs (n = 1,933) were identified between C and P plants, even though most cold-induced DEGs return to baseline expression levels 10 days after treatment (Fig. 1b), suggesting longer-term maintenance of the DNA methylation changes in cold-treated plants. Consistent with the above results, most cold-induced DMRs were found in the CHH context (Fig. 2c). CHH DMRs were most prevalent in intergenic and promoter regions, while were also overlapped with protein-coding genes, TEs, and ncRNAs (Fig. 2c, Fig. S3a). We identified relatively few DMRs that were associated with DEGs within their gene bodies or promoter regions (Fig. S3b–c). Taken together, the results demonstrate that cold stress induces DNA methylation changes in both gene and TE sequences in the *Arabidopsis* genome. Moreover, the enhanced CHH hypomethylation in the PT condition compared to the T condition in gene bodies and TE (Fig. 2a) suggests that cold priming induced by the first stress may lead to an amplified epigenomic response upon exposure to a second cold treatment.

### Cold stress memory genes are marked with lower DNA methylation and an open chromatin state

To investigate the role of DNA methylation in the regulation of cold stress memory genes, we profiled methylation levels in the gene body and ±2 kb flanking regions of cold stress memory genes (n = 500). Cold memory genes exhibited consistently lower DNA methylation levels across all three cytosine contexts compared to randomly selected genes (Fig. 3a–b). CG methylation levels in the memory genes were consistently lower than in random genes across all conditions (Fig. 3b). Similarly, CHH methylation was significantly lower in the memory genes under PT treatment compared to randomly selected genes. In addition, significant demethylation of cold memory genes was observed in the PT group relative to C and T, further confirming that cold priming induces substantial DNA hypomethylation (Fig. 2a, 3b). These results indicate that cold stress memory genes are generally hypomethylated across all sequence contexts and undergo enhanced CHG and CHH demethylation in response to repeated cold stress.

**Figure 3.**
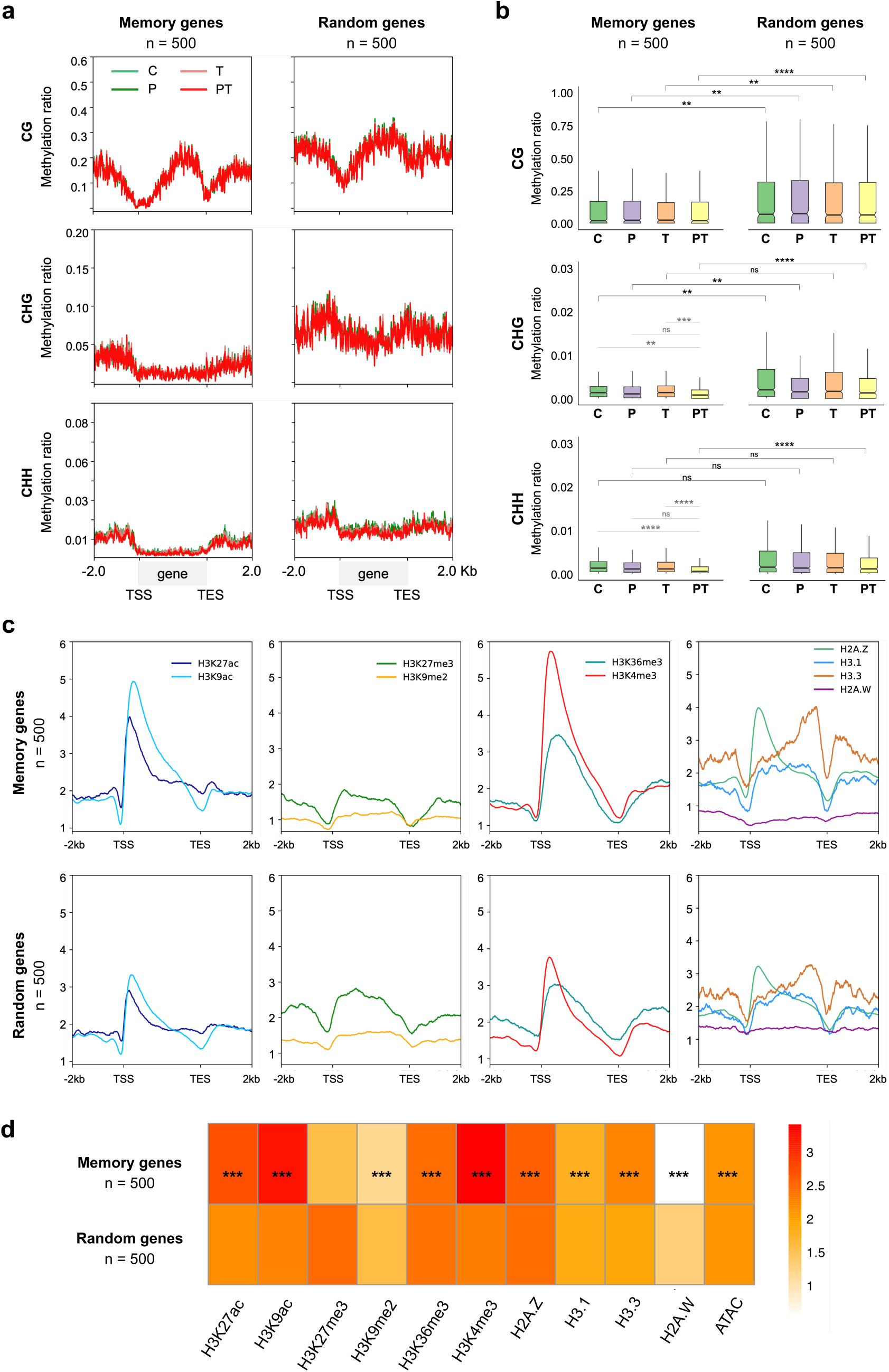
DNA methylation and chromatin states in cold stress memory genes. **a.** Metaplots of DNA methylation level over gene bodies and the 2.0 kb up- and down-stream regions of the WT cold memory genes (n = 500) and randomly selected genes (n = 500). Cytosines with a read coverage greater than 3 were used in the analysis. TSS: transcription start site; TES: transcription end site. **b**. Boxplots of mean gene methylation in cold stress memory genes and random genes in the CG, CHG, and CHH contexts. BH-corrected P-values were calculated using Wilcoxon rank-sum tests (ns: not significant, *p* > 0.05; *, *p* ≤ 0.05; **, *p* < 0.01; ***, *p* < 0.001, ****, *p* < 0.0001). **c**. Metaplots showing enrichment of histone modifications and histone variants over gene bodies and 2.0 kb up- and down-stream regions of the WT cold stress memory genes and random genes. ChIP-seq data was downloaded from the Plant Chromatin State Database (PCSD) ^50^. **d**. Heatmap of enrichment for histone marks between WT cold memory genes and randomly selected genes (n = 500). Statistical significance was assessed using Wilcoxon rank-sum tests (ns: not significant; *, *p* ≤ 0.05; **, *p* < 0.01; ***, *p* < 0.001) with BH adjustment of P-values.

DNA methylation is known to interact with other histone marks and variants to regulate chromatin accessibility, gene expression, and activity of TEs ^49^. Thus, transcriptional activity of genes is often defined by chromatin states – combinations of histone modifications, DNA methylation, and chromatin accessibility. Among the thirty-six chromatin states defined by Liu *et al*. (2018) ^50^, WT cold memory genes were significantly enriched in chromatin states 1 – 4, which are characterized by the actively transcribed chromatin marked by the H3.3 histone variant (Fig. S4a, Supplementary Table 2). Additionally, cold memory genes significantly overlapped with chromatin states 6 and 7, which are marked by the active histone modification H3K36me3. Chromatin states 17, 18, and 20 – 28, described as accessible DNA enriched with active histone marks, such as histone acetylation, H3K4me2, and H3K4me3, were also enriched among the cold memory genes – suggesting these genes reside in a generally open chromatin state. Similar enrichment patterns were observed using chromatin states, defined by Sequeira-Mendes *et al*. (2014) ^51^ (Fig. S4b, Supplementary Table 2).

To validate these findings, we obtained publicly available ChIP-seq data for histone acetylation (H3K9ac, H3K27ac), histone methylation (H3K4me3, H3K36me3, H3K27me3, H3K9me2), histone variants (H2A.Z, H3.1, H3.3. H2A.W), and ATAC-seq DNA accessibility data from the PCSD ^50^. Compared to randomly selected genes, cold memory genes showed strong enrichment of H3K9ac and H3K27ac around the transcription start site (TSS) and within gene bodies, while also exhibiting significantly higher levels of the active histone marks H3K4me3 and H3K36me3 (Fig. 3c–d). Additionally, a significantly lower level of the repressive mark H3K9me2 was observed within and around gene bodies of the memory genes (Fig. 3c, d, Supplementary Table 3). Cold memory genes also displayed slightly but significantly higher levels of the histone variants H2A.Z and H3.3, while being depleted of the repressive variant H2A.W (Fig. 3d, Supplementary Table 3). Finally, cold memory genes were significantly enriched for ATAC-seq signal (Fig. 3d, Supplementary Table 3), strongly suggesting that these genes are generally associated with open chromatin states.

Within the cold memory genes, Cluster I (n = 220) – defined by low basal expression and upregulation upon cold treatment – showed higher CG body methylation than Cluster II (n = 264) genes, which are highly expressed under control conditions and downregulated by cold stress (Fig. S5), although overall epigenomic features were similar. Nevertheless, the gene bodies of Cluster I were more enriched for H3K36me3 but exhibited lower levels of histone acetylation and DNA accessibility (Fig. S5b, c, Supplementary Table 3).

### DNA methylation mutants acquire a wider set of cold stress memory genes

Since cold stress memory genes showed differential DNA methylation profiles compared to other genes in WT plants, we hypothesized that cold-induced gene priming might be affected in mutants impaired in DNA methylation. To test this hypothesis, we analyzed the transcriptomic response of the cold primed and/or triggered *drm2*, *met1*, *ddm1*, and *cmt2/3* mutant plants. The global transcriptomic profile of the mutants closely resembled those of WT plants (Fig. S1a, Fig. S6). Interestingly, however, the WT cold memory genes were not differentially expressed between primed and non-primed plants in *met1* (Fig. 4a, Fig. S7a) and showed altered responses in other mutants (Fig. S7a–d).

**Figure 4.**
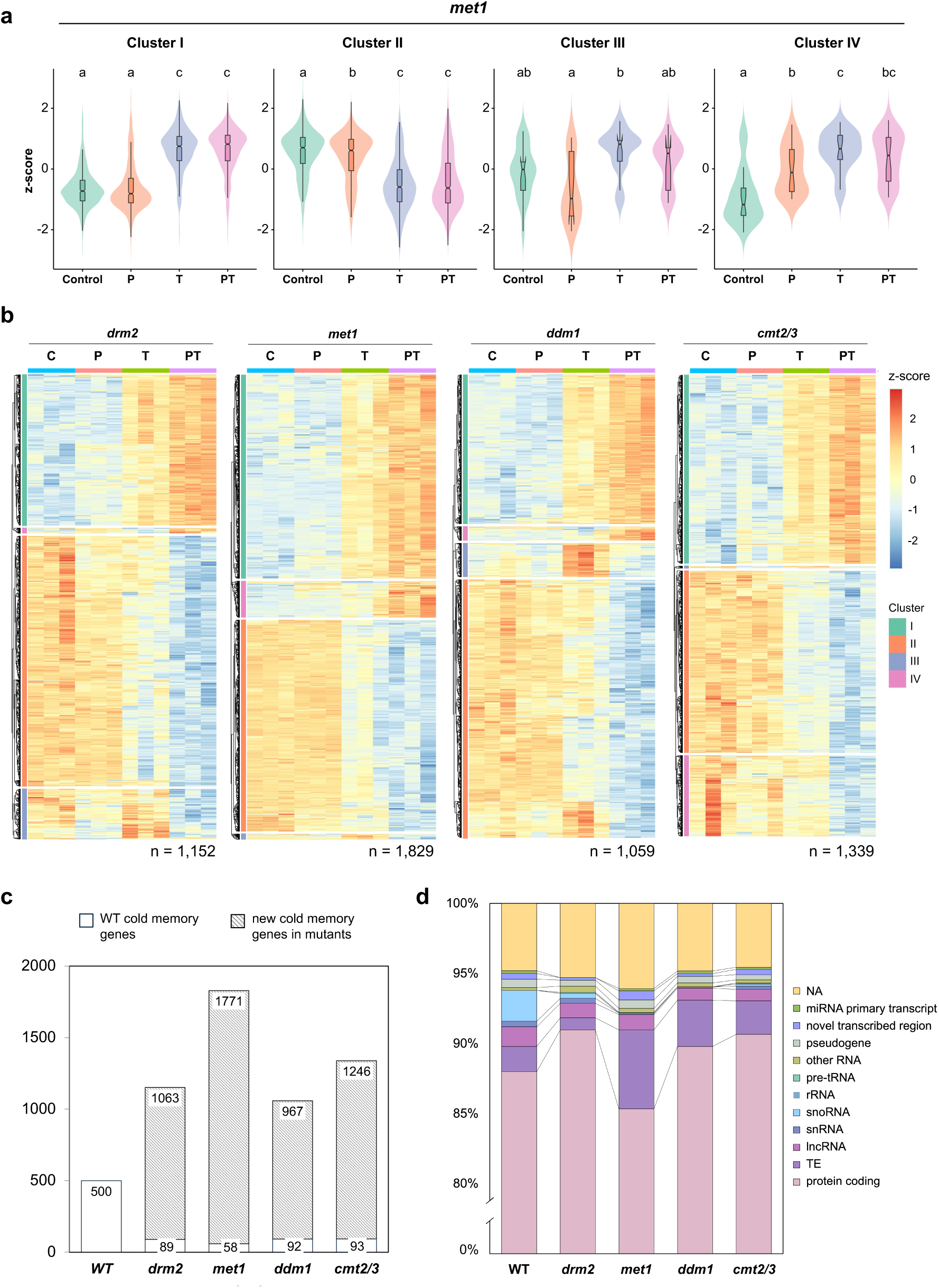
Cold stress memory genes are redistributed in DNA methylation mutants. **a.** Violin plots of mean genes expression of the WT cold stress memory genes in *met1* mutant plants across gene expression clusters. Statistical significance was determined by pairwise Wilcoxon rank-sum tests, with BH method multiple hypothesis testing correction. Different letters correspond to significant differences between the treatment groups (*p-adj* < 0.01). **b**. Heatmaps showing expression of cold memory genes in epigenetic mutants. DEGs were selected according to *p-adj* < 0.05 and absolute value of log2FC > 0.585. Gene expression values (log2FC) were scaled by row to obtain z-scores and clustered using Euclidian distance. **c**. Numbers of cold memory genes in the DNA methylation mutants that are common with the WT cold memory genes (n = 500) and genes that acquired priming in *drm2*, *met1*, *ddm1*, and *cmt2/3*. **d**. Categories of cold memory genes in the WT and DNA methylation mutants. Genes were annotated with AtRTD3 and Araport11 genome annotation. TE: transposable element; lncRNA: long non-coding RNA; snRNA: small nuclear RNA; snoRNA: small nucleolar RNA; rRNA: ribosomal RNA; tRNA: transfer RNA; miRNA: microRNA; NA: newly annotated genes in AtRTD3 (not included in Araport11 annotation).

We then extracted cold memory genes in each mutant using the same criteria applied to the WT transcriptome data. This analysis revealed more than a thousand novel memory genes in each mutant background – a total of 1,829 cold memory genes in *met1*, 1,152 in *drm2*, 1,059 and 1,339 in *ddm1* and *cmt2/3,* respectively (Fig. 4b, c, Fig. S8a–d). However, only a subset of the WT cold memory genes retained their priming-associated expression pattern in the mutants (Fig. 4c), and the majority of cold memory genes were not shared among the mutants (Fig. S9a). Interestingly, despite the globally hypomethylated background of the mutants, cold memory genes in the mutants exhibited reduced DNA methylation levels-particularly in non-CG contexts-similar to those observed in WT plants (Fig. S10). In contrast to cold memory genes, more than 75% of cold-responsive genes were shared between WT plants and at least one mutant background, with 847 cold-responsive genes commonly identified across all five backgrounds (Fig. S9b). These results highlight the role of DNA methylation in regulating cold-primed gene expression in *A. thaliana*. Notably, the number of TEs among cold memory genes was higher in *met1*, *ddm1*, and *cmt2/3* mutants than in WT plants (Fig. 4d), suggesting that derepressed TEs in DNA methylation mutants can become targets of priming under repeated cold stress.

### Cold stress affects gene transcription of DNA methylation modifiers

We examined whether cold stress-induced DNA hypomethylation is due to altered expression of genes involved in the regulation of DNA methylation (Fig. 5a). We found that non-CG methyltransferase gene *CMT2* was significantly downregulated in both T and PT plants compared to the control (Fig. 5a). Moreover, expression of several RdDM pathway regulators, such as *CLSY1*, *SHH1*, *DCL3*, *AGO4*, and *NRPE1*, was also significantly lower in cold-stressed plants (Fig. 5b). Given the role of *CMT2* and the RdDM pathway in maintaining DNA methylation in the CHH context, reduced expression of these genes under cold stress might contribute to the observed CHH hypomethylation in gene bodies and in the cold stress memory genes.

**Figure 5.**
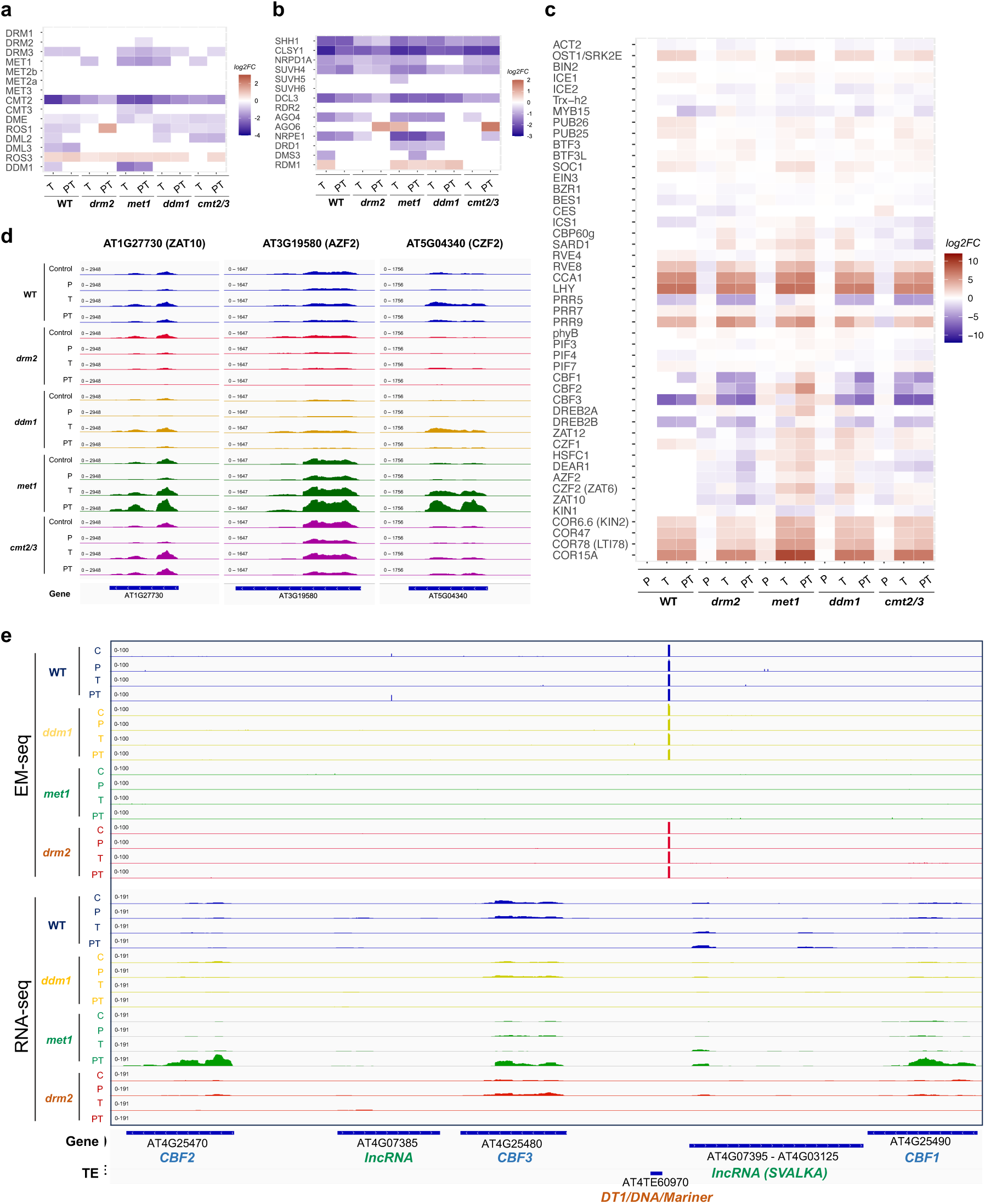
Cold stress and priming responses of epigenetic regulators in *A. thaliana*. **a**, **b**. Heatmaps of expression of the DNA methyltransferases (a) and RdDM pathway regulators (b) in cold-treated WT plants and *drm2*, *met1*, *ddm1*, and *cmt2/3* mutants. Shown are relative expression levels of genes compared to the untreated Control in the corresponding genetic background. The color bar indicates log2FC values obtained from DESeq2 (*p-adj* < 0.05). **c**. Heatmap of the expression of cold stress regulators in cold-treated WT, *drm2*, *met1*, *ddm1*, and *cmt2/3* plants. **d**. Genome browser view of the expression of selected cold response genes. Tracks represent the combined RNA-seq signal from three biological replicates. **e**. Genome browser view of the *CBF* genes showing both gene expression and DNA methylation levels. TE: transposable element.

### Loss of CG methylation in *met1* deregulates the *CBF*-mediated cold response pathway

The *ICE–CBF–COR* cascade plays a central role in plant cold stress responses and is known to be regulated by epigenetic mechanisms, particularly histone modifications ^35,37^. While the role of DNA methylation in this pathway remains less defined, cold-induced methylation changes have been reported across multiple species, including bok choy, tomato, orange, and rubber tree ^52–56^. To understand the role of DNA methylation in the cold stress response pathway, we analysed the expression of common cold-responsive genes in *A. thaliana*. We observed strong upregulation of the *CBF1*, *CBF2*, and *CBF3* genes in the *met1* mutant (Fig. 5c). Similarly, downstream genes – *CZF1*, *AZF2*, *ZAT10*, *COR15A*, and *COR78* – were upregulated in both primed and non-primed plants (Fig. 5c, d). Significant upregulation of these genes was also observed in the *ddm1* mutant under T conditions (Fig. 5c). The expression changes were further validated by RT-qPCR (Fig. S11). In contrast, expression of upstream regulators of the *CBF* genes (*ICE1*, *BTF3s*, *CCA1*, *LHY*, *RVE4/8*) and *ICE1* regulators (*OST1*, *BIN2*) remained unchanged (Fig. 5c).

The expression of *CBF1* under cold stress is tightly regulated by transcription of the antisense lncRNA *SVALKA* ^42^. In addition, transcription at the *SVALKA* locus recruits the PRC2 to the *CBF3* gene, facilitating the deposition of the repressive mark H3K27me3 and downregulating the gene ^43,57^, although the role of DNA methylation in this mechanism remains unclear. We observed a loss of DNA methylation upstream of the *SVALKA* locus in the *met1* mutant (Fig. 5e), while no notable changes in DNA methylation were observed at *CBF* genes between control and cold-treated WT plants (Fig. 5e). Collectively, these findings indicate that *CBF* genes and their downstream targets are specifically deregulated in the *met1* mutant, highlighting a potential role of DNA methylation in the epigenetic control of the *CBF*-mediated cold response pathway.

We examined CG methylation levels at the locus using whole-genome bisulfite sequencing data from 702 Arabidopsis natural accessions ^58^ and found a bimodal distribution of CG methylation (Fig. S12a, b; Table S6). Most accessions carried highly methylated alleles, whereas a small subset (29/702; 4.1%) possessed hypomethylated alleles (mCG < 0.05). Environmental variable data from AraCLIM v2.0 ^59^ suggested that accessions with hypomethylated *CBF* epi-alleles tend to originate from habitats characterized by higher evapotranspiration and greater diurnal temperature range (Fig. S12c, d). We further tested whether the methylation status of the *CBF* locus influences the transcriptional responses of CBF genes and *SVALKA* to cold stress and priming. Under our experimental conditions, however, we detected no clear differences in cold-induced expression changes between hyper-methylated accessions (Col-0, Vid-1, Vim-0, Cvi-0) and hypo-methylated accessions (Lecho-1, Kly-1, Shigu-1) (Fig. S12e–j; Table S7). Thus, the functional contribution of DNA methylation to the CBF-mediated cold response pathway needs further investigation.

## Discussion

In this study, we explored the role of DNA methylation in cold stress priming and memory in *A. thaliana* using transcriptome and methylome analyses of WT and DNA methylation mutants. Cold stress significantly altered expression of thousands of genes, including canonical cold-responsive genes (*COR15A*, *COR47*, *COR78*) (Fig 1, Supplementary Data 1). While the transcriptomic profiles of primed and non-primed plants overlapped considerably, we identified 500 genes differentially expressed only in primed plants, which we defined as cold stress memory genes (Fig. 1c, d). GO analysis revealed that genes related to cold acclimation were generally upregulated in cold-treated plants. In contrast, cold stress memory genes upregulated in the primed plants were associated with metabolic processes and nucleic acid biosynthesis. Nevertheless, both cold-responsive and cold memory genes were enriched for RNA processing and ribosome biogenesis-related processes, confirming the importance of ribosome biogenesis in cold tolerance in plants ^39–41^.

In addition to the protein-coding genes among the cold memory genes, we also identified ncRNAs, including lncRNAs, snRNAs, and snoRNAs, as well as TEs (Fig. 1g). Previous studies have shown that cold stress alters the expression and alternative splicing of 135 lncRNAs, including precursors of microRNAs (miRNAs) and trans-acting small interfering RNAs (ta-siRNAs) ^60,61^. Furthermore, cold-induced lncRNAs, such as *SVALKA*, can confer cold tolerance to plants by regulating the *CBF* genes ^43,62^. Our findings expand on this by showing that cold priming also modulates the expression of various ncRNAs, which include lncRNAs, snRNAs, and snoRNAs (Fig. 1g, Fig. S1f–i). Given the role of priming in enhancing stress tolerance, these results point to a potential regulatory function of ncRNAs in cold adaptation. Additionally, we identified novel TEs including *ATCOPIA46* family that were differentially expressed under cold treatment and priming (Fig. S2). Considering that many TEs are known to influence stress-responsive gene expression and contribute to plant stress resilience ^63–68^, these TEs may act as regulatory elements in cold stress responses in *A. thaliana*.

Genome-wide DNA methylation analysis in primed and non-primed *A. thaliana* plants under cold stress revealed significant hypomethylation in the body regions of protein-coding genes and TEs in the CHH context in cold-triggered plants, with primed plants showing even lower methylation levels than non-primed ones (Fig. 2a–b), suggesting a potential role of DNA hypomethylation in enhancing cold tolerance. This idea is supported by prior studies showing that inhibition of DNA methylation improves freezing tolerance in *A. thaliana* ^69^, and similar cold-induced demethylation has been reported in maize, sugar beet, and chickpea ^70–72^. Correspondingly, DMRs identified between control and cold-treated plants (T and PT) were predominantly hypomethylated and enriched in the CHH context, with over 10% overlapping gene bodies and more than 35% located in promoters (Fig. 2d). Similarly, we observed a higher number of cold-induced DEGs overlapping with promoter DMRs than with gene body DMRs, suggesting that promoter demethylation may play a more prominent role in transcriptional regulation under cold stress.

Interestingly, while the majority of the cold-responsive genes were shared between the DNA methylation mutants and WT plants, less than 10% of the cold memory genes overlapped with those in WT plants (Fig. 4c, Fig. S7a–b). Instead, more than 1,000 new genes were primed in the *drm2*, *met1*, and *cmt2/3* mutants. These findings suggest that DNA methylation plays a greater role in cold priming than in the general cold response, highlighting the epigenetic plasticity of stress memory and its role in restricting spurious gene activation. Comparison of the methylation levels between cold memory genes and random genes revealed that memory genes were significantly hypomethylated in both WT plants and DNA methylation mutants (Fig. 3a, b, Fig. S9), reinforcing the link between low methylation and transcriptional memory. These memory genes were also enriched in chromatin states associated with active transcription, including reduced DNA methylation, active histone modifications, and the H3.3 and H2A.Z histone variants (Fig. 3c–e). The presence of H2A.Z is particularly intriguing, given its known antagonism with CG methylation ^73^ and its association with transcriptionally active, euchromatic regions ^74^. Earlier studies in *A. thaliana* suggested that cold stress treatment promotes acetylation of the histone H3, promoting CBF binding to *COR* gene promoters ^75^. Collectively, these findings suggest a model in which cold stress triggers extensive chromatin reprogramming, with DNA demethylation contributing to the formation of an open chromatin state in primed genes, enabling sustained gene activation via removal of repressive epigenetic marks and enrichment of activating features such as histone acetylation and H2A.Z. Further investigation is needed to understand how cold and priming affect histone modifications and histone variant deposition at the memory genes.

To investigate the mechanism underlying cold-induced DNA hypomethylation, we examined the expression of key DNA methylation regulators. Surprisingly, the DNA demethylases *ROS1*, *DME*, and *DML3* were downregulated in both primed and non-primed plants (Fig. 5a), suggesting that active demethylation is not the primary driver of methylation loss under cold stress. In contrast, we observed significant downregulation of the DNA methyltransferase *CMT2* and several components of the RdDM pathway, including *SHH1*, *CLSY1*, and *NRPD1A* (Fig. 5a–b), indicating that passive demethylation via suppression of methylation maintenance machinery is likely to contribute to the observed DNA hypomethylation in cold-treated plants. These findings are consistent with reports in rubber trees where cold stress induced the downregulation of *CMT* and *DRM* genes ^52^. Additionally, a study in sugar beet reported cold-induced CHH demethylation accompanied by reduced expression of *CMT2* and RdDM-related genes ^71^. Since CMT2 acts in a feedback loop with H3K9 methylation, future work should assess histone methylation dynamics under cold stress. Finally, an assessment of the functional consequences of altered DNA methylation suggested that loss of methylation at the tandem *CBF* genes in *met1* mutants correlated with their overexpression in cold primed plants (Fig. 5c–e), along with that of downstream targets, suggesting a regulatory role for DNA methylation in the *CBF–COR* pathway.

In conclusion, cold priming induced significant transcriptomic changes accompanied by long-lasting DNA methylation changes. The reduced DNA methylation in cold stress memory genes corresponded with epigenetic marks of active transcription, suggesting that it may enable their rapid reactivation. Overall, this work highlights DNA methylation as a key modulator of cold stress memory in plants and provides a foundation for future studies on epigenetic priming mechanisms in crop resilience.

## MATERIAL AND METHODS

### Plant Material

*Arabidopsis thaliana* ecotype Columbia (Col-0) was used as WT plants. The transgenic lines *met1-3*, *drm2-2*, and *ddm1-1*, used in this study were described previously ^76–79^. *cmt2/3* mutant was generated by crossing *cmt3-11* ^79^ and *cmt2-3* (SALK_012874). Second-generation homozygous plants of *met1-3*, *ddm1*, *drm2,* and *cmt2/3* plants homozygous for at least two generations were used in the study. Seeds were germinated and grown in 0.5x Murashige and Skoog (MS) media with 0.8% agar and 0.1% sucrose at 22°C with a long day (16 h light / 8 h dark). Nine days post germination, plants were transferred to soil and kept under the same conditions. For the cold stress experiment, three biological replicates were used for each treatment group. Cold stress was performed as indicated in Figure 1a. In short, after transferring to the soil, the control group (C) was grown at 22°C for 14 days and samples were collected at 25 days. For the priming cold stress treatment, 12-day-old plants were transferred to 4°C for three days. Afterward, plants were transferred to the standard conditions for 7 days. Triggering cold stress was applied to the 22-day-old plants at 4°C for three days. For all groups, leaf samples were collected at 25 days post germination, frozen in liquid nitrogen and stored at - 80°C.

### RNA extraction and sequencing

For RNA extraction, about 100 mg of frozen samples was homogenized with Micro Smash tissue homogenizer (MS-100R) (TOMY, Japan) and total RNA was extracted with the Maxwell® 16 LEV Plant RNA Kit (Promega Corporation, USA) following instructions from the manufacturer. The purity of RNA was assessed by nanodrop spectrophotometer NanoDrop One^C^ (Thermo Fisher Scientific Inc., USA) and samples were stored at -80°C.

For RNA-sequencing, 1 μg of total RNA was quantified with Qubit™ RNA Broad Range (BR) Assay Kit (Thermo Fisher Scientific (Lifetechnologies), USA). Next, rRNA depletion was performed using the QIAseq FastSelect –rRNA Plant Kit (QIAGEN, Germany) and libraries were prepared using NEBNext Ultra II Directional RNA Library Prep Kit for Illumina (New England Biolabs, USA). Library quality and average read length were assessed using the Agilent 2200 TapeStation system (D1000 ScreenTape System) (Agilent Technologies, Germany). For library pooling, the concentration of each library was measured using Qubit 1X dsDNA HS Assay Kit (Thermo Fisher Scientific (Lifetechnologies), USA) and pooled using Illumina Pooling calculator (https://support.illumina.com/help/pooling-calculator/pooling-calculator.htm). Paired-end sequencing of the libraries was performed at Okinawa Institute of Science and Technology Graduate University (OIST) Sequencing Section (SQC) on NovaSeq 6000 (Illumina) (read length 150 bp) (Supplementary Table 5).

### Bioinformatics analysis of RNA-seq data

Raw sequencing reads from RNA-seq were analyzed using the nf-core pipeline “rnaseq” (revision 3.10.1) ^80^. Quality check on the reads was performed with FastQC (v. 0.11.9), and reads were trimmed using Trim Galore! software (v. 0.6.7) (https://github.com/FelixKrueger/TrimGalore). Reads were then mapped to the TAIR10 reference genome with STAR (v. 2.7.9a) ^81^ and quantified with Salmon (v. 1.9.0) ^82^. AtRTD3 transcriptome ^83^ was used for read annotation.

Differential gene expression analysis for RNA-seq was performed in R (version 4.4) using DESeq2 (v. 1.44.0) ^84^. Genes were classified as differentially expressed (DEGs) if the Benjamini-Hochberg corrected *p-adjusted* (*p-adj*) value was less than 0.05. To select for high confidence cold responsive and cold stress memory genes, a threshold of 0.585 was established for the absolute value of log2 Fold Change (log2FC). PCA graphs were plotted with *plotPCA()* function of DESeq2 and with ggplot2 (v. 3.5.1)^85^ using variance stabilizing transformation (vst)-transformed count data. The scale factor for each sample was calculated using *sizeFactors()* in DESeq2 and BigWig (bw) files were created using the *bamCoverage* command of *deepTools* suite (v. 3.4.3) ^86^ with parameters *--binSize 1 --scaleFactor --exactScaling*. Gene tracks were visualized using Integrative Genomics Viewer (IGV) (v. 2.16.2)^87^ and heatmaps were plotted with R packages pheatmap (v. 1.0.12) (https://github.com/raivokolde/pheatmap) and ggplot2 (v. 3.5.1) ^85^. Venn diagrams were plotted using InteractiVenn ^88^. Gene Ontology (GO) analysis was performed with PANTHER^89^.

### Reverse Transcription Quantitative Polymerase Chain Reaction (RT-qPCR)

RT-qPCR was performed using the RNA extracted as described above. cDNA was transcribed using Prime Script™ II 1st Strand cDNA Synthesis Kit (Takara, Japan) according to the manufacturer’s instructions, starting with 1 μg of total RNA. The concentration of obtained cDNA was measured with Nanodrop and qPCR was performed with gene-specific primers using TB Green™ Premix Ex Taq™ II (Tli RNaseH Plus) (Takara, Japan) in the Thermal Cycler Dice ® Real Time System III (Takara, Japan). The specificity of primers was checked using the dissociation curve after the PCR reaction. PCR program included one cycle of initiation at 90°C for 30 sec, followed by 40 cycles of denaturation at 95°C for 5 sec and extension at 60°C for 30 sec. Primers for RT-qPCR were designed manually and/or with QuantPrime ^90^ and are listed in Supplementary Table 4. Gene expression levels were calculated using the 2^−ΔΔCT^ method ^91^ and normalized with *ACTIN* 2 (AT3G18780) and GLYCERALDEHYDE-3-PHOSPHATE DEHYDROGENASE (*GAPDH*, AT1G13440). Statistical differences between gene expression levels were calculated using Student’s t-test at α ≤ 0.05.

### DNA extraction and Enzymatic Methyl sequencing library preparation

Plant material for EM-seq was sampled as described above. Genomic DNA (gDNA) was extracted from around 0.1g of frozen plant tissue following the Nucleon PhytoPure DNA extraction kit (Cytiva, USA) protocol manual. Isolated gDNA was quantified with Qubit dsDNA HS Assay Kit and 50 ng of gDNA was sheared into ∼300 bp fragments using Covaris Focused-ultrasonicator M220 (Covaris, USA). Libraries were constructed using NEBNext Enzymatic Methyl-seq (EM-seq™) kit for Illumina (New England BioLabs, USA) following the manufacturer’s recommendations. NEBNext® Multiplex Oligos for Enzymatic Methyl-seq (Unique Dual Index Primer Pairs) (New England BioLabs, USA) were used for multiplexing the libraries. Libraries were quantified using the Qubit dsDNA HS Assay Kit and the average size of the libraries was determined with Agilent 2200 TapeStation system. Paired-end (150 bp) sequencing was carried out at the OIST SQC section using the Illumina NovaSeq6000 sequencer (Supplementary Table 5).

### Bioinformatics analysis of EM-seq data

For reproducible processing of the raw sequencing reads from EM-seq, nf-core pipeline “*methylseq*” (revision 2.5.0) was used with the following parameters: *--comprehensive true --em_seq true --cytosine_report true*. Read quality control and trimming were performed with FastQC (v. 0.11.9) and Trim Galore! (v. 0.6.7), respectively. BAM files from replicates were merged and reads were then aligned to the TAIR10 genome. Methylation calls were extracted using the Bismark (v. 0.24.0) software. Alignment quality was assessed with Qualimap (v. 2.2.2-dev). Following the read processing with the nf-core pipeline, cytosine reports (CX_report) were manually generated using the *bismark_methylation_extractor* function of Bismark (v. 0.23.0) with options *--CX --cytosine_report*. Methylation levels of the chloroplast DNA were used to estimate the rate of the enzymatic conversion (higher than 99%). Metaplots were plotted using *computeMatrix* function of *deepTools* suite with parameters *-bs 10 -p max/2* and using the *plotProfile* function with *--perGroup* parameter. Differentially methylated regions (DMRs) were identified using the DMRcaller software (1.30.0) ^92^ with *computeDMRs* function. Following parameters were used: *regions = NULL, method = “bins”, windowSize = 100, kernelFunction = “triangular”, test = “fisher”, pValueThreshold = 0.05, minCytosinesCount = 4, minProportionDifference = 0.1, minGap = 200, minSize = 50, minReadsPerCytosine = 4, cores = 1*. Cutoff of 30%, 20%, and 10% difference was used for the DMRs in the CG, CHG, and CHH contexts, respectively. Genes overlapping with the DMRs were extracted using the *bedtools intersect* command (v. 2.92.2) ^93^. To calculate mean methylation levels, only cytosines with a minimum coverage of 3 were selected. Violin plots and boxplots were plotted in R using *ggplot2*.

### Chromatin states analysis

For chromatin states’ analysis, genomic regions with defined chromatin states were downloaded from Liu et al., ^50^ and ^51^. Cold stress memory genes and all genes in the TAIR10 genome were then overlapped with the genomic regions in a given chromatin state. The command line tool, *bedtools intersect* with option *-wo* was used to identify regions with at least 150 bp overlap, which approximately corresponds to the length of DNA wrapped around one nucleosome. Enrichment for a chromatin state within the cold stress memory genes was tested with Fisher’s exact test (*p-adj* < 0.01).

### Natural accession analysis

Processed whole genome methylome data from 972 *Arabidopsis* natural accession were obtained from 1001 Genome Project ^58^. From this dataset, CG DNA methylation levels at the CBF locus (Chr4: 13,020,033–13,020,046; comprising 8 CG sites) were extracted using a minimum read coverage threshold of >=3 reads per cytosine. This filtering resulted in valid methylation profiles for 702 accessions. Environmental variable data were obtained from AraCLIM v2.0 ^59^. The variables were standardized by Z-score transformation. To identify environmental factors associated with methylation variation, we categorized accessions into two groups based on average CG methylation across the 8 CG sites in the CBF locus: High-methylation accessions: mean CG methylation > 0.9 (n = 518); Low-methylation accessions mean CG methylation < 0.05 (n = 29). PCA was applied to the Z-score–normalized environmental variables, and t-tests were conducted for each variable to compare the two groups. Multiple testing correction was performed using the False Discovery Rate (FDR) method, with a significance threshold of FDR < 0.05.

## Data availability

The sequencing data generated in this study have been deposited in EMB-EBI European Nucleotide Archive database under accession code PRJEB92364. ChIP-seq data for histone acetylation (H3K9ac – SRX648276, H3K27ac – SRX1469074), histone methylation (H3K4me3 – SRX1518741, H3K36me3 – SRX1518744, H3K27me3 – SRX853860, H3K9me2 – SRX361927), histone variants (H2A.Z – SRX1436239, H3.1 – SRX130311, H3.3 – SRX130301, H2A.W – SRX352374), and ATAC-seq data (ATAC – SRX2000799) were obtained from the Plant Chromatin State Database (PCSD) ^50^.

## Acknowledgement

We thank the OIST SQC for RNA-seq and EM-seq sequencing services. We thank Dr. Leonardo Furci and Dr. Jérémy Berthelier for assistance in performing the cold stress experiment and for their useful input during project discussions.

