## Supplemental Figures for "Transcriptome and epigenome dynamics underpin cold stress priming in *Arabidopsis*"

### Supplementary Figures

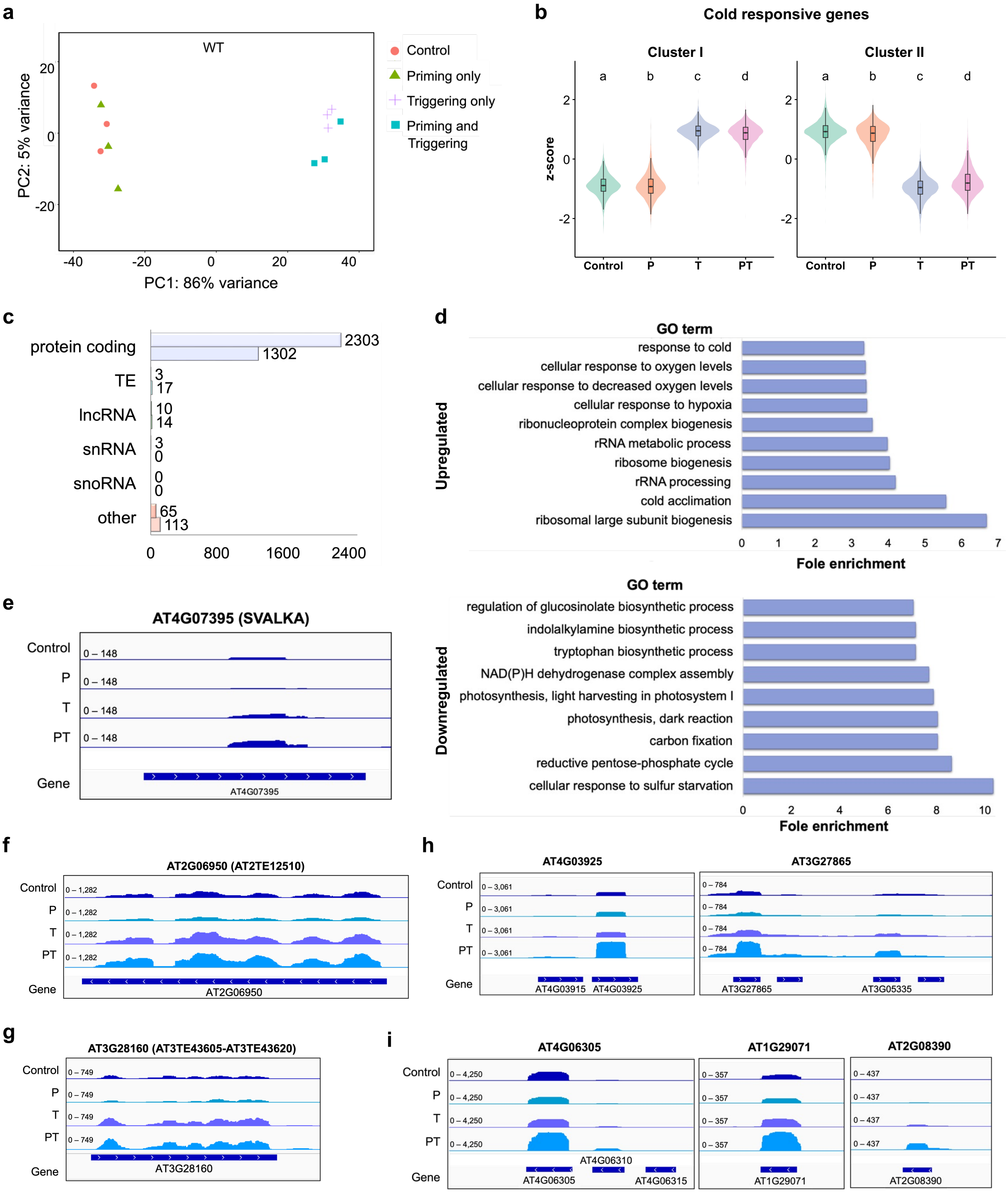


**Supplementary Figure S1. Cold stress priming alters transcriptomic response in *A. thaliana.***

**a**. Principal Component Analysis (PCA) plot for WT plants treated with Priming only (P), Triggering only (T), Priming and Triggering (PT), or no cold stress treatment (Control, C). The data scores after variance-stabilizing transformation (vst) are shown (n = 3). **b**. Violin plots of mean expression of cold-responsive genes clusters in WT. Statistical significance was determined by pairwise Wilcoxon rank-sum tests, with Benjamini-Hochberg (BH) multiple testing correction. Different letters indicate significant differences between the treatment groups (*p-adj* < 0.01). **c.** Gene categories among the cold-responsive genes in WT (n = 3,830). Top bar: genes upregulated in PT; bottom bar: genes downregulated in PT plants. pre-tRNA, ribosomal RNA, miRNA primary transcript, pseudogene, and novel transcribed region were classified as 'other'. **d**. GO term analysis of cold-responsive genes in WT. Fold enrichment of GO terms with Bonferroni-corrected P-values < 0.05 are displayed. **e-i**. Genome browser view of cold-responsive lncRNA *SVALKA* (e), cold memory TEs (f-g), snRNAs (h) and snoRNAs (i). Tracks represent the combined RNA-seq signal from three biological replicates.

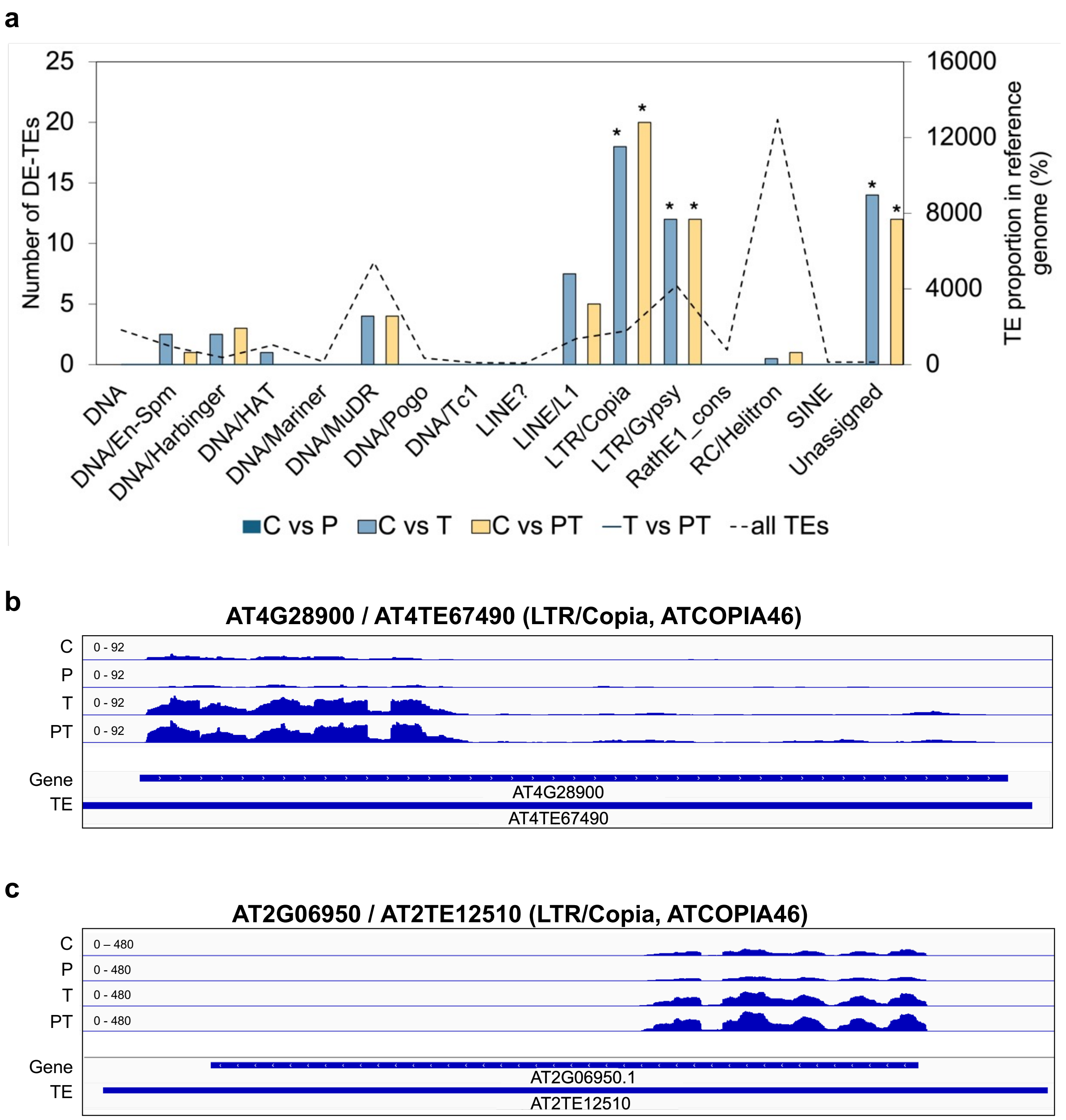


**Supplementary Figure S2. Differentially expressed TEs (DE-TEs) in cold-treated plants**.

**a**. Enrichment for TE families among DE-TEs in WT plants. The dashed line represents proportions of each TE superfamily in the genome of *A. thaliana*. Bars represent the number of DE-TEs. Enrichment was tested using Fisher's exact test ($p < 0.05$). **b**. Genome browser view of cold-responsive LTR/*Copia* superfamily TEs. Tracks represent the combined RNA-seq signal from three biological replicates.

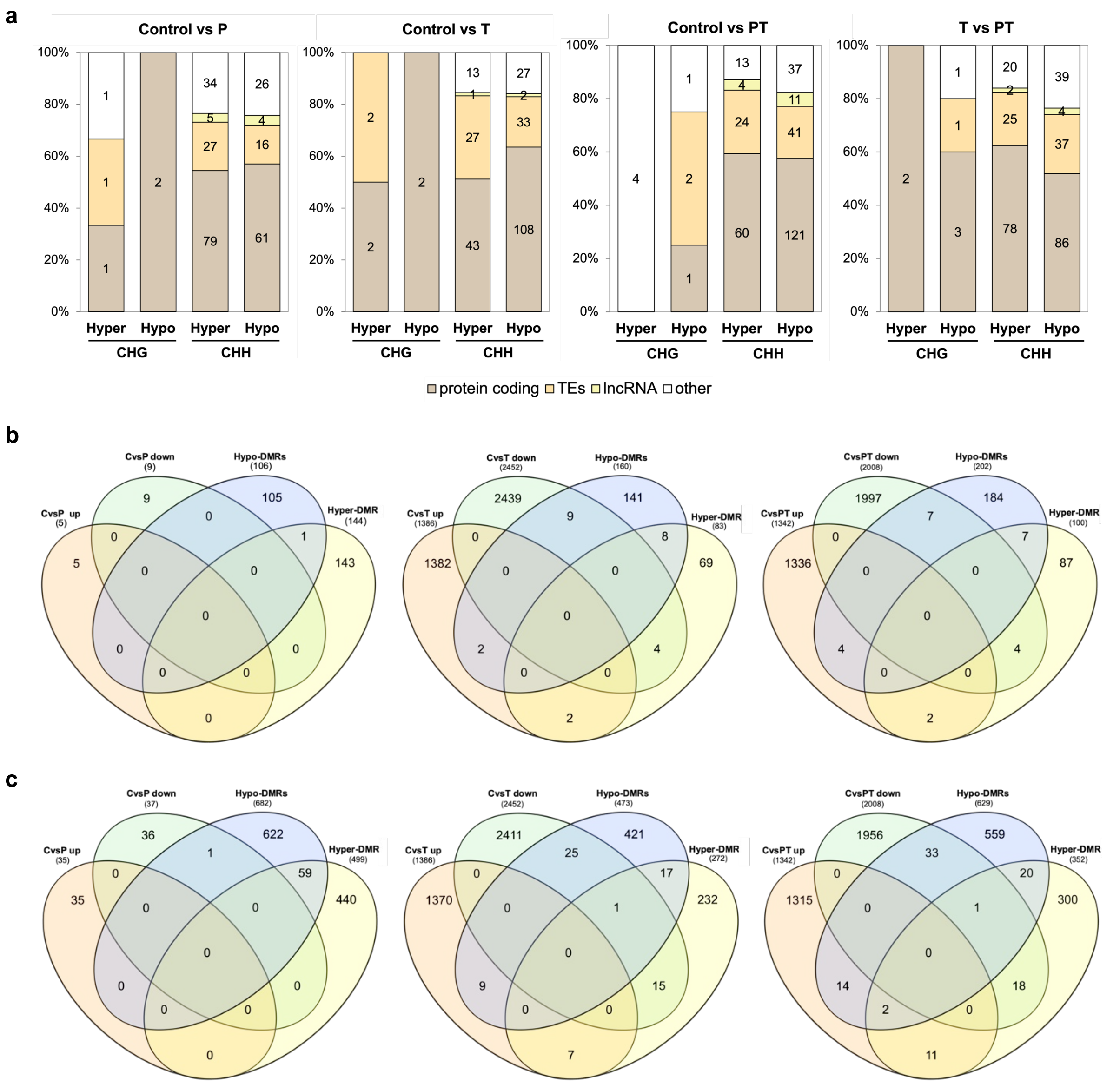


**Supplementary Figure S3. DNA methylation associated with differentially expressed genes in cold-treated plants.**

**a**. Number of DMRs in protein-coding genes, TEs, and ncRNAs. Genes were classified using Araport11 and AtRTD3 annotations. TEs: transposable elements; lncRNA: long-non-coding RNA; other: novel transcribed regions, pseudogenes, other RNA, and novel genes annotated in AtRTD3, but not Araport11. **b-c**. Venn diagrams showing the overlap between DEGs (*p-adj* < 0.05 and |log2FC| > 0.585) and DMRs in gene bodies (b) and promoters (c) in the corresponding comparison: Control vs. P, Control vs. T, and Control vs. PT. Up: upregulated genes; Down: downregulated genes; Hypo-DMRs: hypomethylated DMRs; Hyper-DMRs: hypermethylated DMRs. Genes overlapping more than one DMR were counted once.

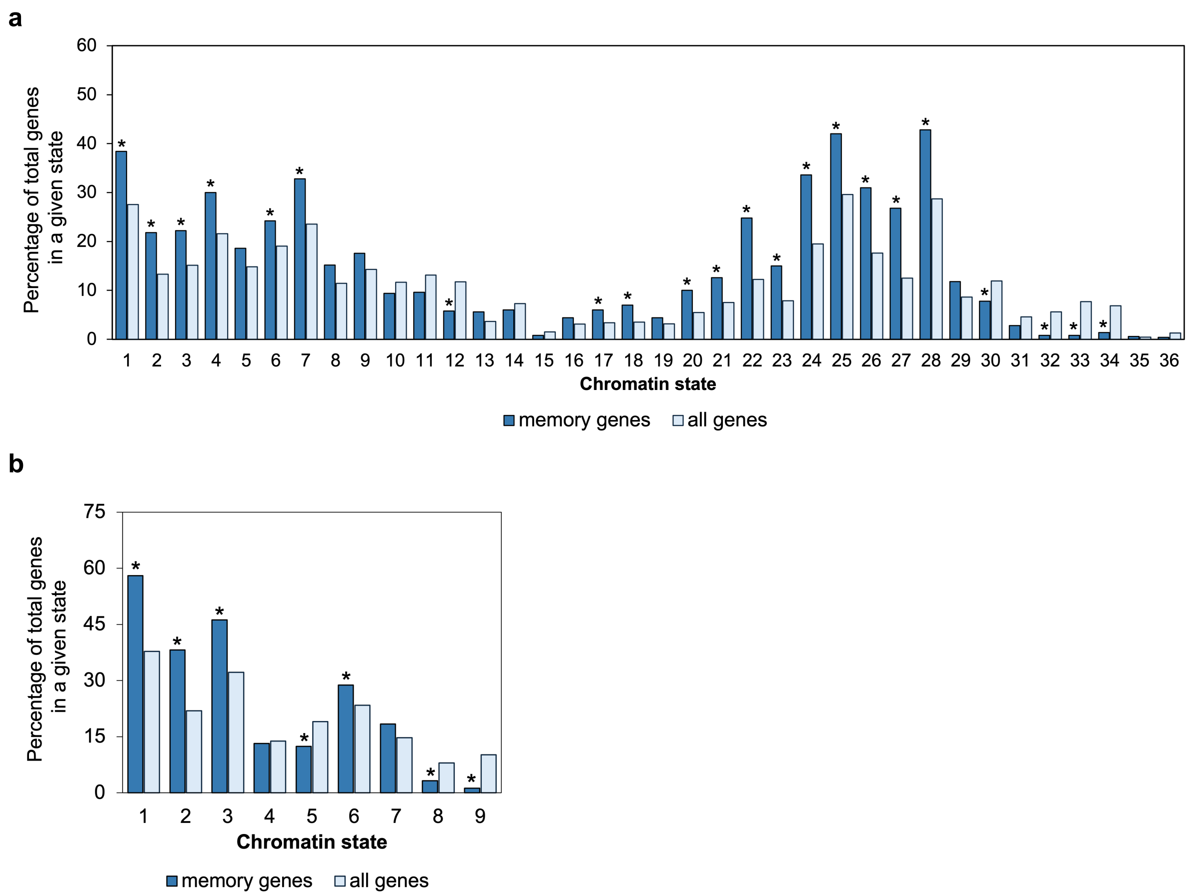


**Supplementary Figure S4. Chromatin state of cold stress memory genes.**

**a-b**. Overlap between WT cold stress memory genes and chromatin states from Liu et al. (2018) (a) and Sequeira-Mendes et al. (2014) (b). Enrichment was tested with Fisher's exact test (*p-adj* < 0.01)**.**

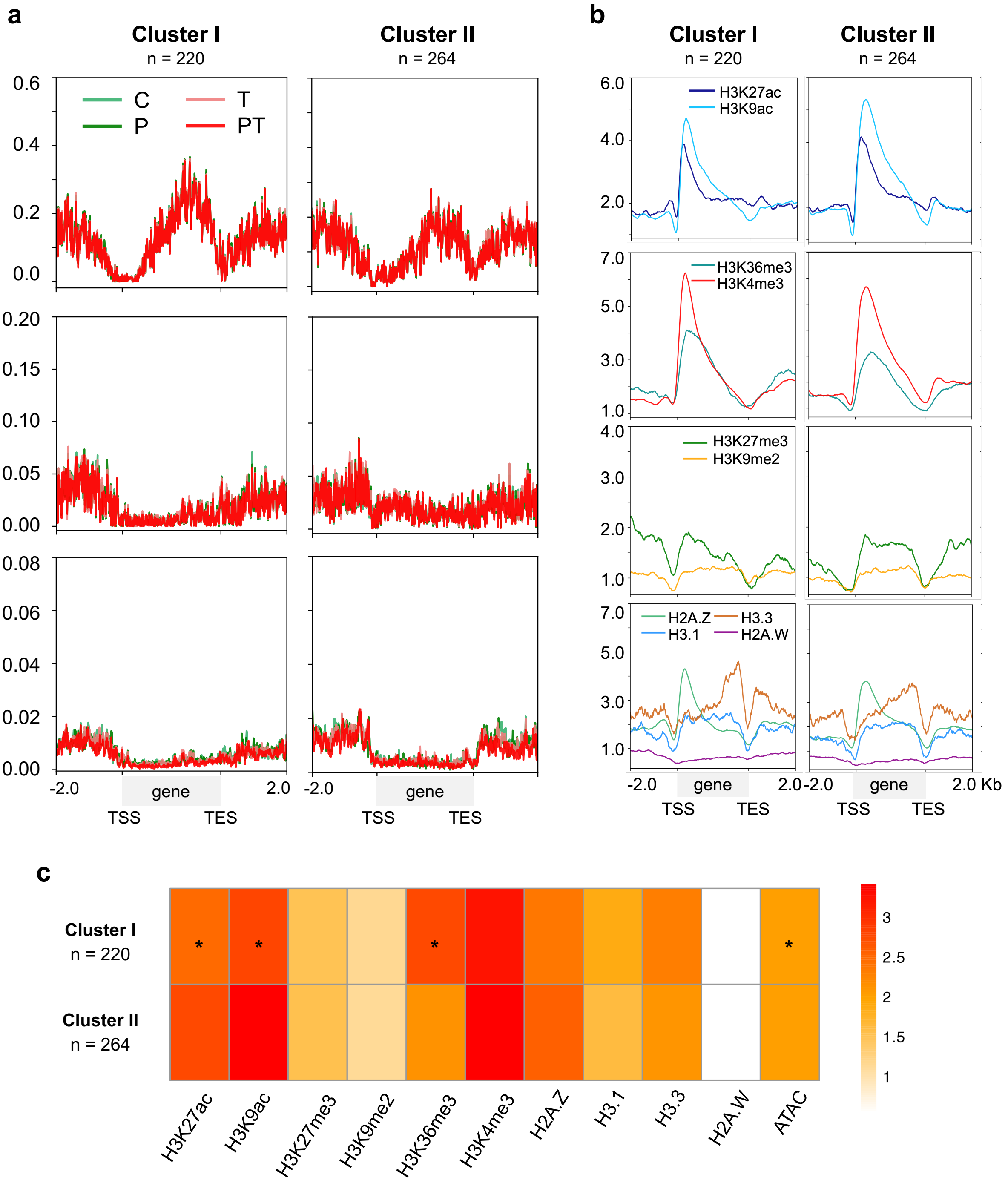


**Supplementary Figure S5. DNA methylation and chromatin states in Cluster I and II cold stress memory genes in WT.**

**a-b.** Metaplots of DNA methylation level (a) and histone modifications and histone variant enrichment (b) over gene bodies and the 2.0 kb up- and down-stream regions of the WT cold memory genes Cluster I (n = 220) and Cluster II (n = 264). Cytosines with read coverage greater than 3 were used in (a). ChIP-seq data (b) was downloaded from the Plant Chromatin State Database (PCSD) [50]. TSS: transcription start site; TES: transcription end site. c. Heatmap of enrichment for histone marks between Clusters I and II of the WT cold memory genes. Statistical significance determined by Wilcoxon rank-sum tests (ns: not significant; *, $p \leq 0.05$; **, $p < 0.01$; ***, $p < 0.001$) with BH adjustment of P-values.

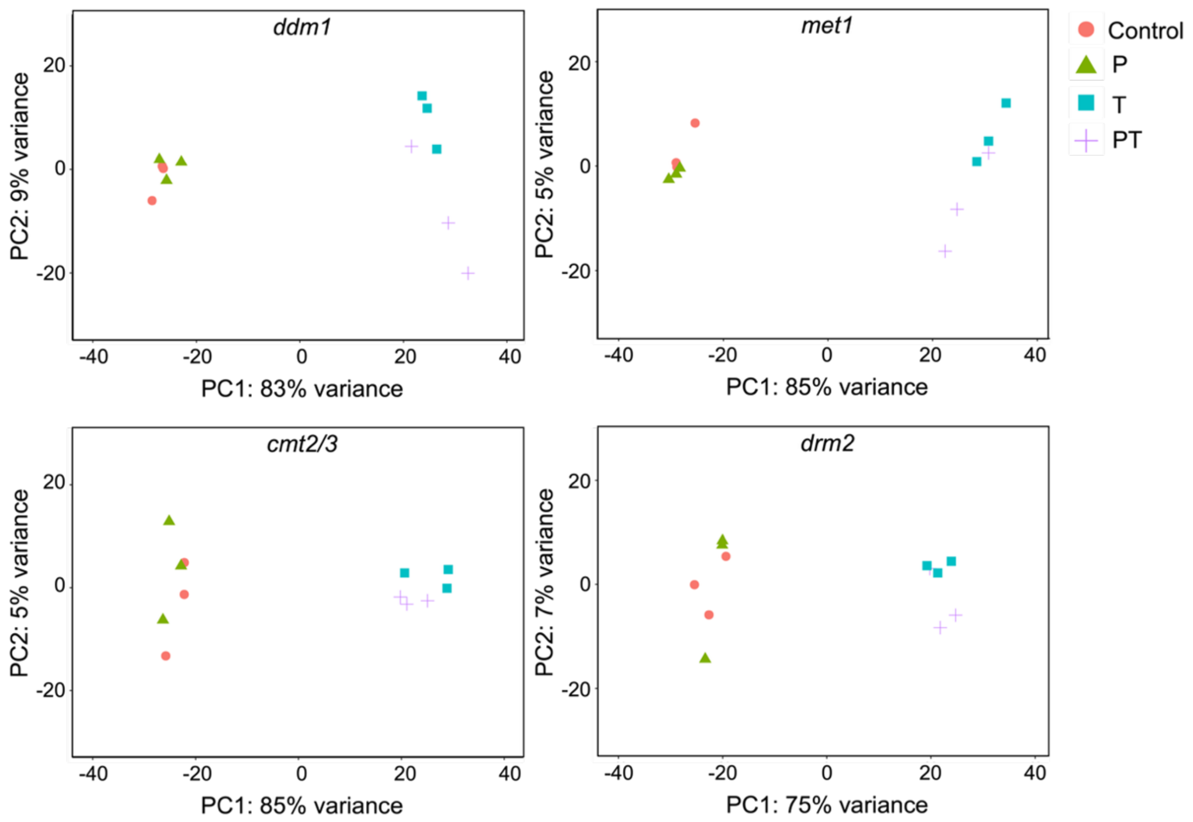


**Supplementary Figure S6. Transcriptomic responses of *Arabidopsis* DNA methylation mutants with cold stress.**

PCA plots for *ddm1*, *met1*, *cmt2/3*, and *drm2*. The plots show PC1 and PC2 for untreated control and cold-treated (P, T, PT) plants. Shown are scores of VST data (n =3).

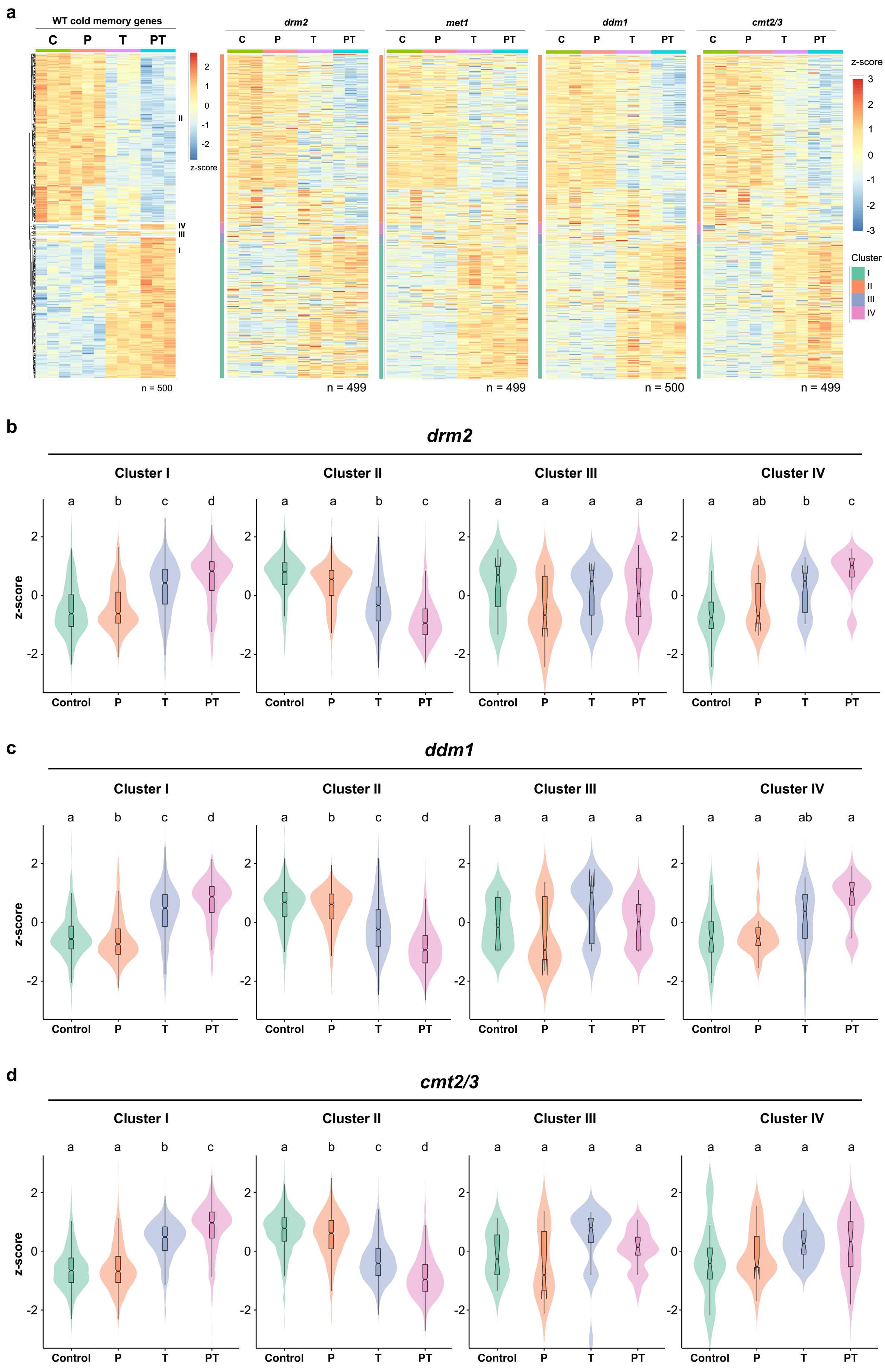
a
WT cold memory genes
C
P
T
PT
II
IV
III
I
z-score
n = 500
drm2
met1
ddm1
cmt2/3
n = 499
n = 499
n = 500
n = 499
Cluster
b
drm2
Cluster I
Cluster II
Cluster III
Cluster IV
z-score
Control
c
ddm1
d
cmt2/3

**Supplementary Figure S7. Changes in transcriptomic responses of cold stress memory genes in DNA methylation mutants.**

**a.** Heatmaps of expression of cold stress memory genes in epigenetic mutants. DEGs were selected according to *p-adj* < 0.05 and absolute value of log2FC > 0.585. Gene expression values (log2FC) were scaled by row to obtain z-scores and clustered using Euclidian distance. **b-d**. Violin plots of mean gene expression of the WT cold memory genes (n= 499~500) in *drm2* (b), *ddm1* (c) and *cmt2/3* (d) mutant plants by clusters. Statistical significance was determined by pairwise Wilcoxon rank-sum tests, with BH multiple testing correction. Different letters correspond to significant differences between the treatment groups (*p-adj* < 0.01).

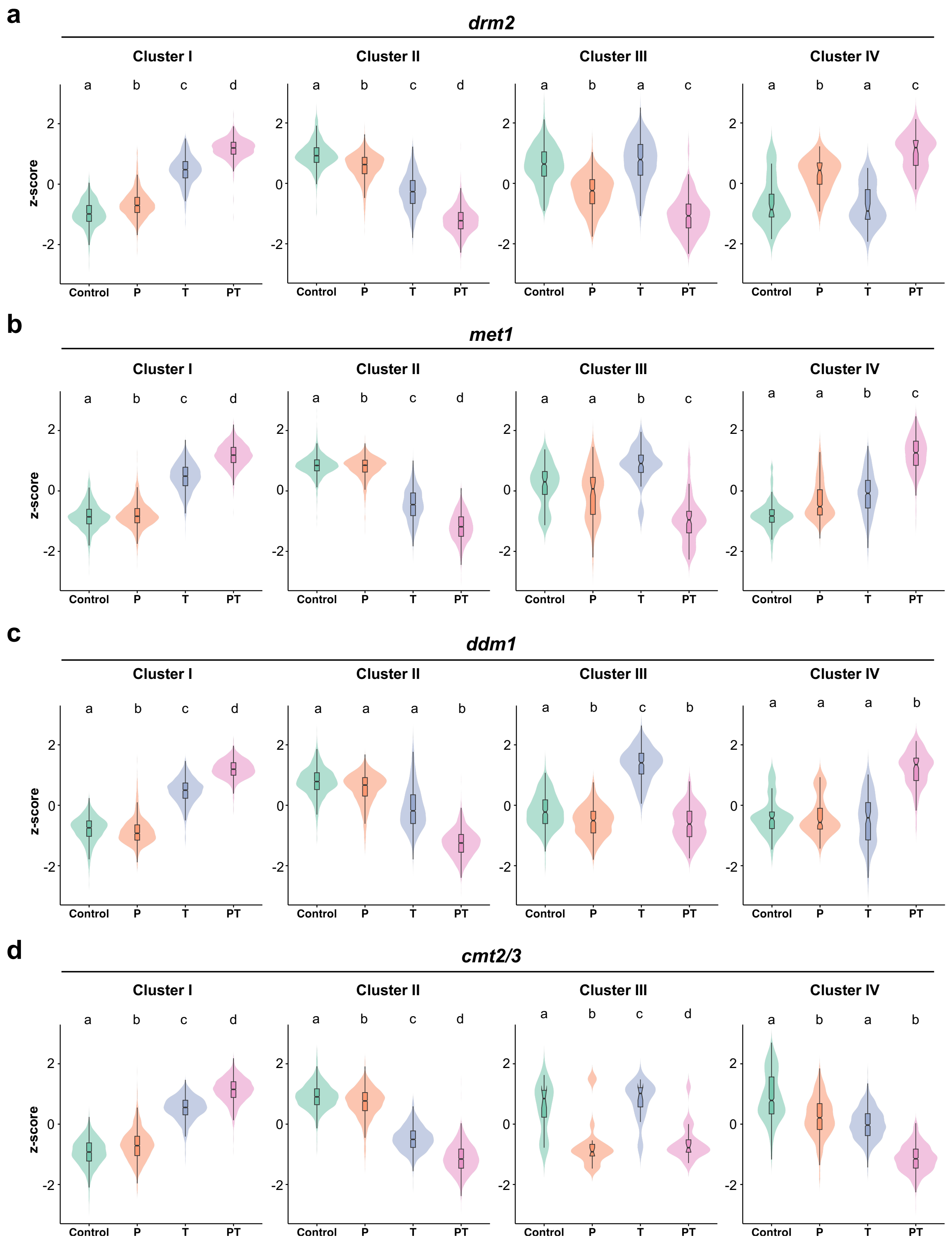


**Supplementary Figure S8. Cold memory genes acquired in DNA methylation mutant.**

**a-d.** Violin plots representing mean z-scores of expressions of genes by cluster for cold stress memory genes identified in *drm2* (a), *met1* (b), *ddm1* (c), and *cmt2/3* (d). Different letters indicate statistically significant differences between the groups ($p < 0.01$). P-values were determined by pairwise Wilcoxon rank-sum tests, with BH multiple testing correction.

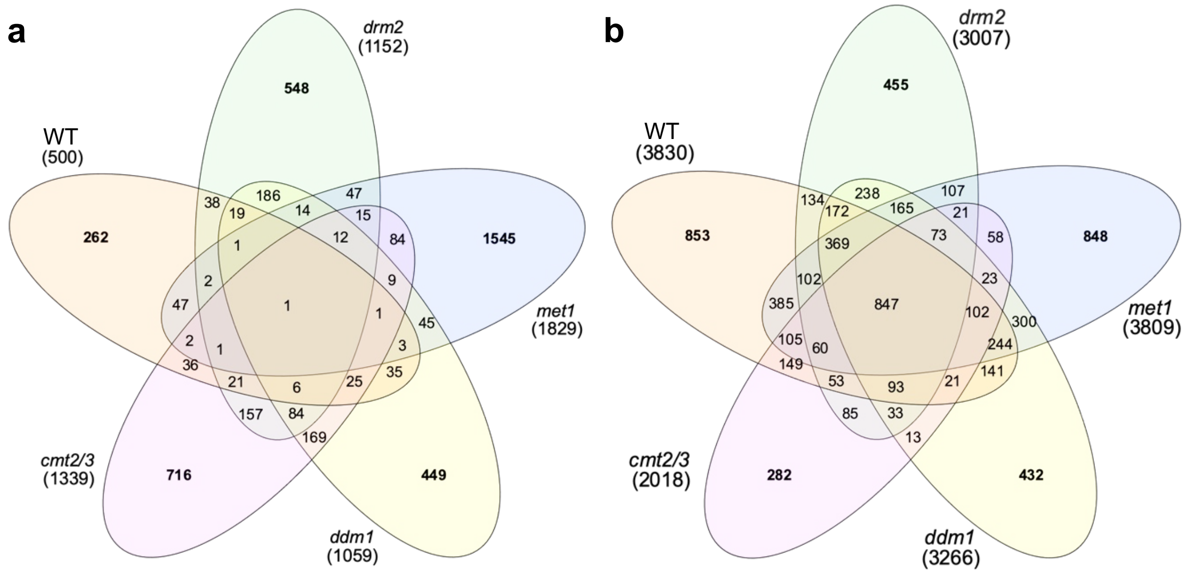


**Supplementary Figure S9. Cold stress memory genes and cold-responsive genes in DNA methylation mutants.**

**a-b**. Venn diagrams for the overlap between cold memory genes (a) and cold-responsive genes (b) in WT and the DNA methylation mutants.

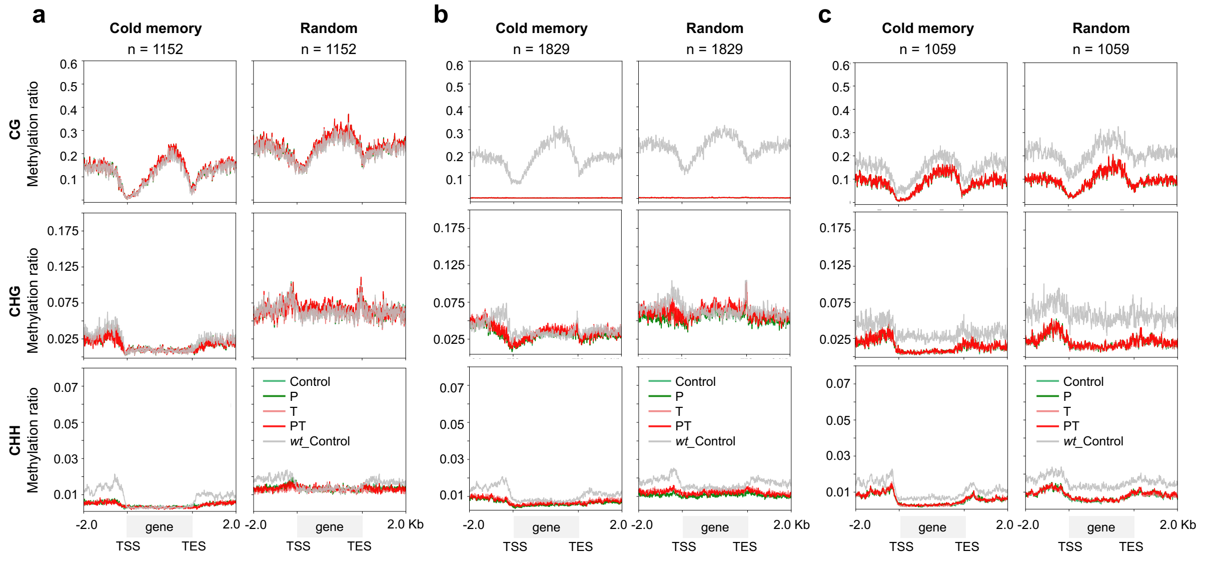


**Supplementary Figure S10. DNA hypomethylation of cold stress memory genes in DNA methylation mutants.**

**a-c.** Metaplots of the cold stress memory genes in mutants in *drm2* (a), *met1* (b), and *ddm1* (c). Metaplots were generated over gene bodies and the 2.0 kb up- and down-stream region of the genes. DNA methylation levels in WT in corresponding genic regions were plotted in gray lines. Cytosine with read coverage greater than 3 were used in the analysis. TSS: transcription start site; TES: transcription end site.

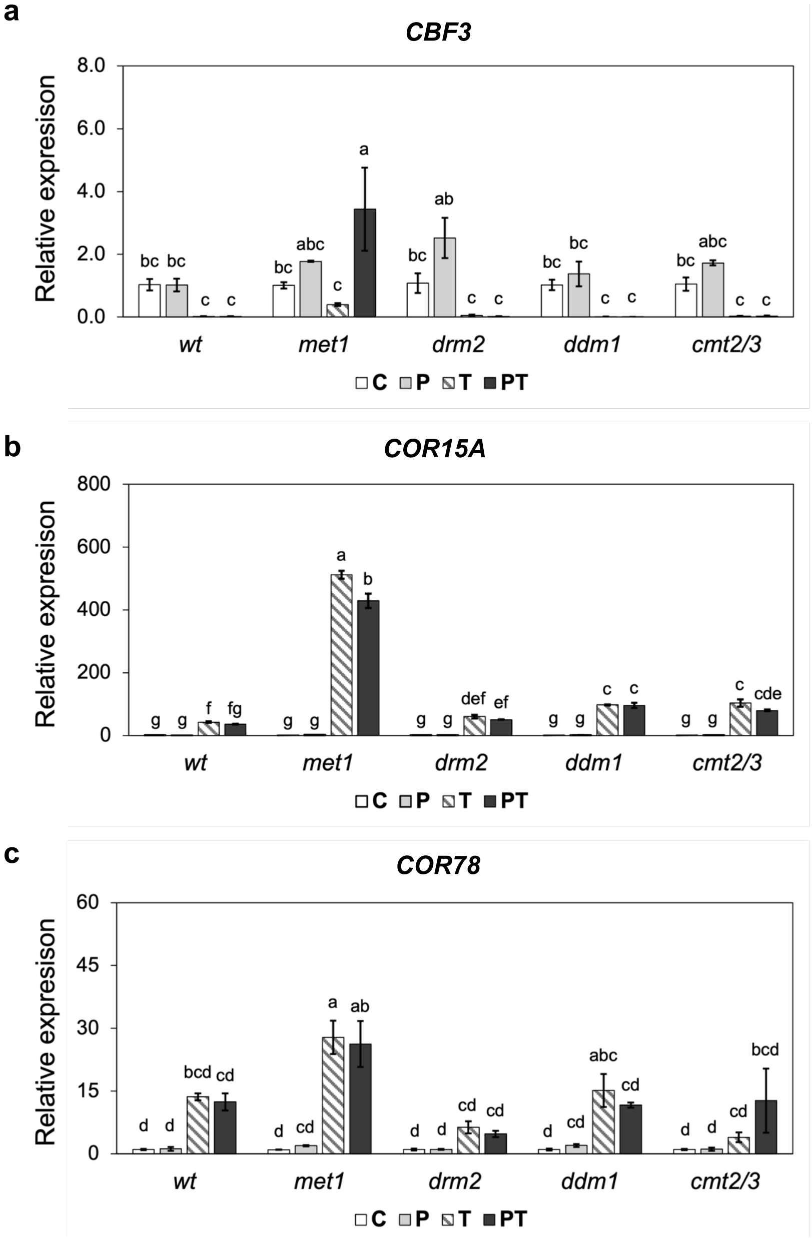


**Supplementary Figure S11. Deregulation of the *CBF*-mediated cold response genes in DNA methylation mutants.**

Relative expression of *CBF3* (a), *COR15A* (b), and *COR78* (c) genes. Relative expression was measured in control and cold-treated plants by RT-qPCR (primers listed in Table S4) and normalized against *ACTIN 2* and *GAPDH*. Bars represent the mean of three replicates ± SD. Different letters indicate statistical significance determined by two-way ANOVA followed by Tukey's HSD test ($p < 0.05$).

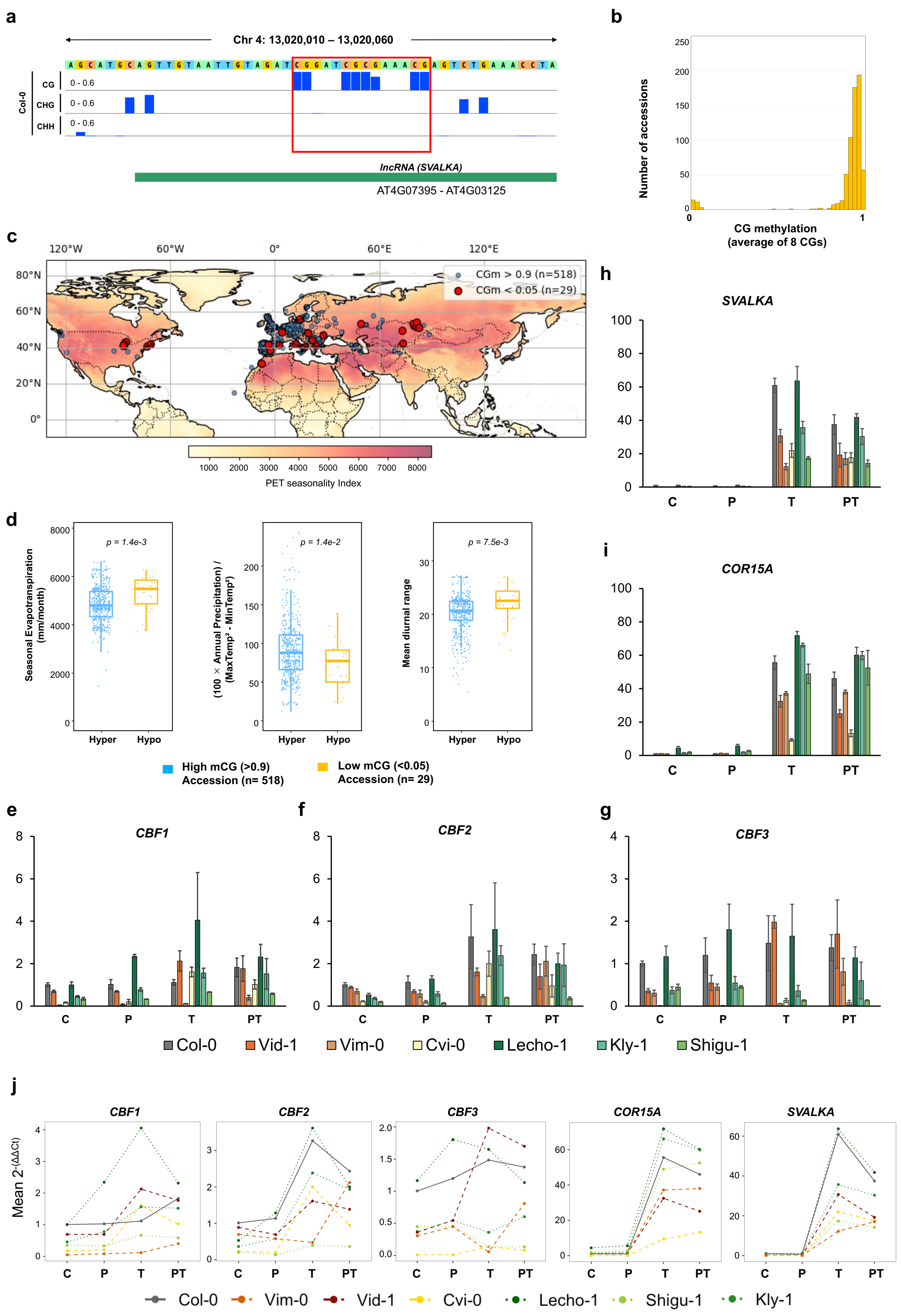

a
Chr 4: 13,020,010 – 13,020,060
Col-0
CG
CHG
CHH
0 - 0.6
lncRNA (SVALKA)
AT4G07395 - AT4G03125
b
Number of accessions
CG methylation
(average of 8 CGs)
c
CGm > 0.9 (n=518)
CGm < 0.05 (n=29)
PET seasonality Index
d
Seasonal Evapotranspiration (mm/month)
p = 1.4e-3
(100 × Annual Precipitation) / (MaxTemp² - MinTemp²)
p = 1.4e-2
Mean diurnal range
p = 7.5e-3
Hyper
Hypo
High mCG (>0.9) Accession (n= 518)
Low mCG (<0.05) Accession (n= 29)
e
CBF1
f
CBF2
g
CBF3
h
SVALKA
i
COR15A
C
P
T
PT
Col-0
Vid-1
Vim-0
Cvi-0
Lecho-1
Kly-1
Shigu-1
j
CBF1
CBF2
CBF3
COR15A
SVALKA
Mean 2^-(ΔΔCt)
Col-0
Vim-0
Vid-1
Cvi-0
Lecho-1
Shigu-1
Kly-1

**Supplementary Figure S12. DNA methylation of the *CBF* locus in *Arabidopsis* natural accessions.**

**a.** CG sites in the *CBF* locus (Chr4: 13,020,033−13,020,046; comprising 8 CG sites) examined in natural accessions. **b**. DNA methylation level of the CG sites in natural accessions (n=702). **c**. Sampling location of natural strains with hyper-methylated CGs (>0.9, dark gray circle) and hypo-methylated CGs (<0.05, red circle) in the *CBF* locus. Potential evapotranspiration (PET) seasonality index is overlayed as heatmap on the world map. **d**. Boxplots of environmental variables obtained from AraCLIM v2.0 (59) are plotted for strains with hyper-methylated CGs and hypo-methylated CGs in the *CBF* locus. P-values were determined by Brunner-Munzel test. **h-j.** RT-qPCR for *CBF1-3* genes, *COR15A*, and *SVALKA*. Relative expression was measured in control and cold-treated plants by RT-qPCR and normalized against *ACTIN 2* and *GAPDH*. Bars represent the mean of three replicates ± SD.
